# Actin network heterogeneity tunes activator-inhibitor dynamics at the cell cortex

**DOI:** 10.1101/2025.07.28.667299

**Authors:** Ondrej Maxian, Aaron R. Dinner, Edwin Munro

## Abstract

Biological systems can display diverse patterns of self-organization, even when built on conserved networks of interaction between molecular species. In these cases, reaction-diffusion equations provide a valuable tool to learn how new dynamics could emerge from quantitative tuning of parameters. Bringing these models into quantitative correspondence with biological data remains an outstanding challenge, especially when the data manifest heterogeneities that are difficult to account for mathematically. One particular example occurs in cell biology, where the membrane-bound regulatory protein RhoA interacts with the filamentous actin cortex in an activator-inhibitor loop. Though this core biochemical circuit is conserved across multiple cell types in different organisms, it produces different patterns of RhoA activity in different contexts, from traveling waves in starfish to transient pulses in *C. elegans*. To understand how this variation emerges, we develop an activator-inhibitor model that accounts explicitly for actin assembly and heterogeneity. By fitting the model to summary statistics of experimental data, subject to known parameter constraints, we show that F-actin assembly dynamics tune the spatiotemporal patterns of RhoA activity. A minimal representation of these dynamics reveals how directional transport (via polymerization) combines with stochasticity in F-actin number and orientation to produce the observed patterns. This work sheds light on how phenotypic diversity arises from heterogeneity and anisotropy, with important implications for the next generation of activator-inhibitor models.

**Significance:** To divide, move, and polarize, cells must self-organize their constituent proteins into large-scale patterns with varied spatiotemporal character. The design principles of this process remain poorly understood, primarily because quantitatively matching mathematical models to experimental data is difficult. Here we consider pattern formation from two constituents on the cell cortex: the regulatory protein RhoA and actin filaments. Using a mathematical model, constrained quantitatively by data from multiple organisms, we show how diversity in RhoA activity can arise from intra- and inter-organismal changes in actin filament architecture and assembly dynamics. Our results reveal general principles for pattern formation at the cortex, and our combination of data analysis, modeling, and parameter inference provides a broadly-applicable, interdisciplinary methodology to unravel mechanisms of self-organization.

## Introduction

In cellular and developmental biology, large-scale patterns emerge from self-organization of smallscale constituents [1]: of molecules within cells [2–5] and cells within tissues [6, 7]. In cells in particular, the structure, function, and interactions of key proteins are often conserved, yet these proteins can self-organize into different structures in different contexts [8,9], implying that variations in emergent patterns must come from quantitative differences in the underlying parameters of interaction. Understanding how molecular-scale parameter tuning contributes to cell- and tissue-scale variation remains challenging, however, both because of the difficulty of making direct parameter measurements *in vivo*, and because developmental patterning involves collective dynamics with complicated feedbacks. Consequently, mathematical models play an essential role in inferring how changes in parameters could produce new phenotypes.

Arguably the most influential class of developmental patterning models are reaction-diffusion models [1,10–14], which clearly demonstrate how changes in parameters (e.g., the diffusion constant of each species) can give rise to dramatically different self-organized spatiotemporal patterns (e.g., standing patterns and traveling waves) [13,15–19]. To date, however, these models have largely been used qualitatively, for instance to explore the emergence of patterns [5, 20–25] or to differentiate mechanistic hypotheses [22, 26] (see [14] for a review), rather than to fit data directly and infer parameter values [27–29]. The latter approach is more difficult because of the trade-off between model and experimental complexity; for robust inference, the model must be complex enough to describe all of the underlying experimental data, but simple enough to make the inverse problem well-posed [30]. Because of this, parameters often represent effective interactions of many molecules, and inferred values cannot be compared directly with experimental measurements for validation [27, 31–34].

In this paper, we study a paradigmatic example of molecular self-organization in cells, the dynamic coupling of the small GTPase RhoA and the cortical actin cytoskeleton which lies just beneath the cell membrane (Fig. 1A). The actin cytoskeleton is a filamentous meshwork, with architecture set by a balance of assembly dynamics and active remodeling [35–43], while RhoA (hereon “Rho”) is a molecular switch which cycles between an active state and inactive state on the membrane [44–47]. Active Rho promotes further activation of Rho through GTP exchange factors (GEFs), and engages downstream effectors, such as formin and profilin, to nucleate, polymerize, and crosslink actin filaments (F-actin) [44, 45, 48]. Once assembled, F-actin recruits inhibitors (GTPase-activating proteins, GAPs) to inactivate Rho [21, 49], closing a negative feedback loop, and completing the activator-inhibitor circuit (Fig. 1A).

**Figure 1:**
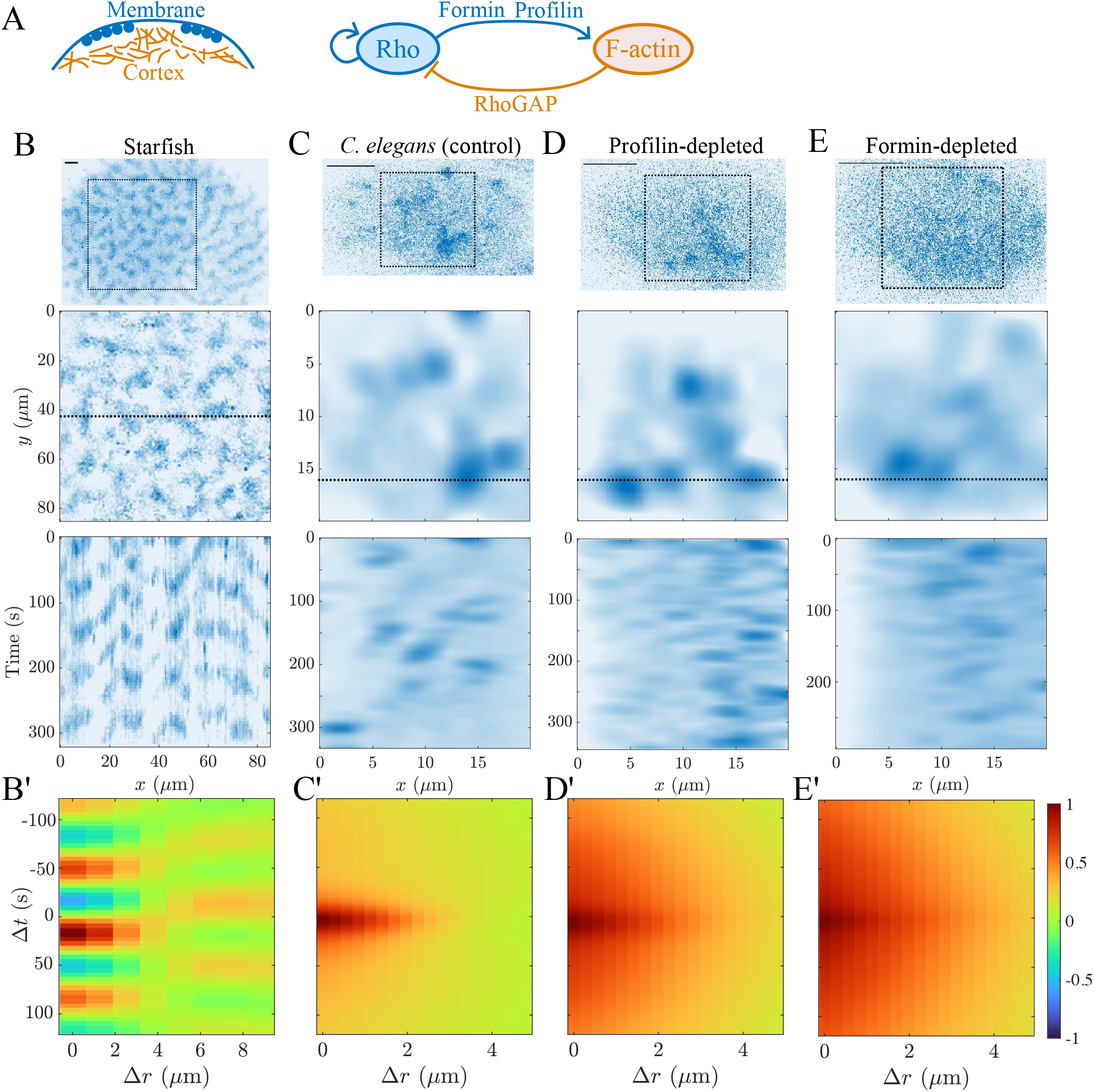
Diverse patterns of RhoA and F-actin are observed *in vivo*. (A) Rho and filamentous actin form an activator-inhibitor system at the cell cortex, with membrane-bound Rho assembling cortical actin filaments through its effectors (including formin and profilin), and actin filaments inhibiting Rho through RhoGAP. (B–E) Rho dynamics in (B) starfish [21, Fig. 1C] and (C–E) *C. elegans* embryos [(C) myosin(RNAi), (D) myosin+profilin(RNAi), and (E) myosin+formin(RNAi) [55]]. Top panels: raw imaging data, scale bars = 10 *µ*m. Middle panels: extracting a square region and filtering the result to identify excitations. Bottom panels: kymographs over the indicated *y*-slice (dotted lines in middle panels). (B’–E’) Cross correlation (1) between Rho and actin for each data set in (B–E).

While the core molecular components of this circuit are broadly conserved across model organisms [20,21,49], the spatiotemporal dynamics of Rho are qualitatively different in different contexts. In frog and echinoderm embryos [20,21,50–53], traveling waves of Rho activity are observed, consistent with theoretical expectations for a rapidly-diffusing activator. In worm embryos, however, the same biochemical circuit produces transient and seemingly-stochastic focal “pulses” of Rho activity [49, 54, 55], which move slowly through the cortex before being extinguished. As we later show, the behavior in worms is not readily reproducible using reaction-diffusion models [56], making it unclear how differences in patterns could arise from manipulation of the biochemical circuit.

Here we show quantitatively that differences in Rho activity in different model systems can be explained by variations in actin network architecture and assembly dynamics. We do so by designing and implementing a “hybrid” model which couples a continuum reaction-diffusion equation for Rho to a discrete actin network undergoing turnover. This model not only reproduces all of the experimentally-observed phenotypes, but also infers kinetic parameters for F-actin turnover that are consistent with available independent measurements, suggesting that it captures key determinants of the variation in dynamics observed *in vivo*. To extract the minimal mechanisms governing this variation, we introduce a continuum model that captures the essential features of F-actin assembly from the hybrid model. This exercise identifies directional transport of inhibition (via F-actin polymerization) and stochasticity in filament number and orientation as principle determinants of variation in Rho dynamics, revealing a key design principle for biological control of pattern variation. Our overall approach integrating model development, data-driven inference, and mechanistic insight has the potential to be broadly applicable to other systems which feature structural diversity [57–60].

## 1 Variation in spatiotemporal Rho dynamics from conserved activator-inhibitor coupling with F-actin

### 1.1 Experimental data reveal context-dependent dynamics

Spatiotemporal dynamics of Rho activity can be read out in different systems by making time-lapse movies of fluorescently-tagged biosensors that selectively bind active Rho. To establish a framework for comparison of patterns across systems, we standardize a set of movies [21, 55] from starfish oocytes and *C. elegans* embryos by selecting a representative sample of the relevant dynamics. To remove boundary effects, we focus on a region in the embryo interior, then perform further filtering to remove noise (see Methods).

In starfish ooctyes, direct observations of a RhoA biosensor reveal a clear pattern of traveling waves [20, 21, 50]. In our representative example (Fig. 1B), these traveling waves are approximately synonymous with locally oscillatory dynamics, which have a period of roughly 60 s (see kymograph in Fig. 1B). By contrast, imaging the Rho biosensor in one-cell *C. elegans* embryos (Fig. 1C) yields a much lower signal-to-noise ratio, with significantly less regularity in the patterns of Rho excitations. Filtered images and kymographs (Fig. 1C) show irregular patches of excitation which seem to appear and disappear stochastically. These excitations have lifespans of about 25 s, during which they meander slowly in space, while maintaining sizes of about 10 *µ*m^2^.

### 1.2 F-actin assembly dynamics modulate Rho activity

Simple activator-inhibitor models predict the traveling waves of Rho excitation observed in starfish oocytes [19–21, 50], but not the pulses observed in *C. elegans*. One notable difference between the systems is that there is significantly more actin network heterogeneity in *C. elegans* embryos [49] than in starfish and frog oocytes [20]. Previous *in silico* studies [56, 61–64] suggest that this heterogeneity could contribute to a wider range of model behaviors. Similarly, previous experimental perturbations showed that modifying F-actin assembly dynamics could induce changes in patterns of Rho activity [55, 65–67]. We thus hypothesize that the actin network architecture and assembly dynamics are responsible for the observed differences in patterns.

As a first step toward testing this hypothesis, we re-examine previously reported [55] spatiotemporal Rho dynamics in *C. elegans* embryos depleted of profilin (which promotes filament elongation [35, 68–70]) or formin (which promotes filament nucleation and elongation [36, 68, 69]). It was previously shown that the essential features of Rho dynamics observed in wild-type cells do not depend on myosin II activity (see Fig. S4 and [55]), while the phenotypes produced by depleting formin or profilin do depend in part on myosin II through the induction of contractile instabilities [55]. Therefore to avoid the confounding effects of myosin II and isolate effects due to variation in actin network architecture and assembly dynamics, we focus here on the analysis of phenotypes produced by depleting formin and profilin in a myosin-mutant background. Compared to control (myosin mutant) embryos, myosin mutant embryos co-depleted of either formin or profilin exhibit larger Rho excitations (Fig. S2–S3). In filtered data, these excitations tend to recur in similar locations more frequently than in control embryos (Fig. 1D and E; see also [55, Fig. 8]). Because formin and profilin mediate F-actin assembly, these data indicate that variation in actin network architecture and assembly dynamics tune the duration and spatial spread of Rho excitations [55].

### 1.3 Statistics for model inference

Given the inherent noise in the Rho activity data, the different systems can only be quantitatively compared statistically. Following previous work [20, 21], we use the cross correlation function

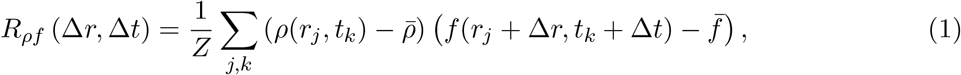

between active Rho (*ρ*; the Rho biosensor) and F-actin (*f* ; the RhoGAP/LifeAct signal) as a means to assess spatiotemporal correlation of the two species. Here 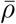 denotes the mean of *ρ* in space and time (likewise for 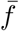), and the normalization constant *Z* is chosen so that max(|*R*|) = 1. Intuitively, (1) denotes the correlation between *ρ* and *f* a distance Δ*r* away and time Δ*t* later. For example, *R*_*ρf*_ (Δ*r* = 0, Δ*t* = −10) < 0 implies a correlation between Rho and a lack of F-actin 10 s earlier, which suggests that Rho spreads into regions that lack F-actin (as is the case for traveling waves).

The cross correlation function puts the contrast between the starfish and *C. elegans* data in sharp relief (Fig. 1B’ and C’): whereas the starfish cross correlation function is oscillatory in space and time in accordance with locally oscillatory traveling waves, the *C. elegans* cross correlation function is positive almost everywhere, with a maximum at (Δ*t*, Δ*r*) = (0, 0) that decays on length (2 *µ*m) and time (25 s) scales that correspond to the typical size and lifetime of an excitation region. The cross correlation functions for the profilin- and formin-depleted embryos are broader in space and time (Fig. 1D’ and E’), consistent with the larger and longer-lived excitations (Fig. S2–S3) in those systems.

## 2 Discrete-continuum model to infer F-actin assembly dynamics from experimental data

To understand how variations in Rho dynamics emerge from changes in F-actin assembly and architecture, we introduce a mathematical model coupling a continuum description of Rho activity to discrete filament dynamics. By forcing the model to reproduce experimental cross correlation functions (under constraints on F-actin assembly), we infer the F-actin dynamics responsible for different patterns of Rho activity.

### 2.1 Coupling discrete actin assembly with continuum Rho dynamics

We use a two-dimensional description of Rho and F-actin dynamics because the membrane and cortex are thin relative to the length of an actin filament [71, 72]. The evolution of active Rho *ρ*(***x***, *t*) is given by [49]

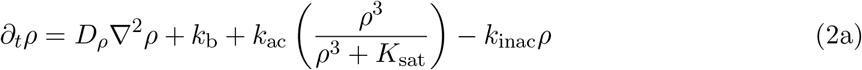

and accounts for basal (*k*_*b*_) and auto-catalytic activation of Rho (through the cubic Hill function). Inactivation of Rho enters through the term *k*_inac_(***x***, *t*), which will later become a function of the F-actin concentration. Since the activation rates (*k*_b_ and *k*_ac_) in (2a) are constant, our model implicitly assumes that inactive Rho is never (substantially) depleted. This assumption is supported by a recent study in starfish [26] which showed that local peaks in Rho activity actually precede peaks in total Rho, implying that recruitment of GAPs/F-actin, rather than depletion of total Rho, terminate local excitation of Rho. In *C. elegans*, the evidence is less direct, but a recent study [73] showed that total Rho exhibits similar spatiotemporal dynamics to active Rho, also suggesting that inactive Rho is not depleted.

To represent a heterogeneous F-actin network with anisotropic growth, we utilize a discrete description in which individual filaments undergo nucleation, elongation and disassembly with kinetics similar to those observed in *C. elegans* embryos [70]. At each point in space, we assume that filaments nucleate stochastically and with random orientation at rate

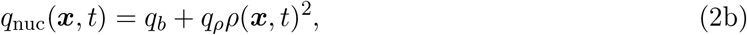

so that nucleation is enhanced in regions of high Rho activity. Once nucleated, a filament polymerizes at the barbed end with rate *ν*_*p*_ to a maximum length *𝓁*_max_, at which it remains fixed for time *T*_fil_, after which it depolymerizes from the pointed end with rate *ν*_*d*_ (Fig. 2A). We allow actin filaments to overlap to mimic the finite thickness of the cortex [71].

**Figure 2:**
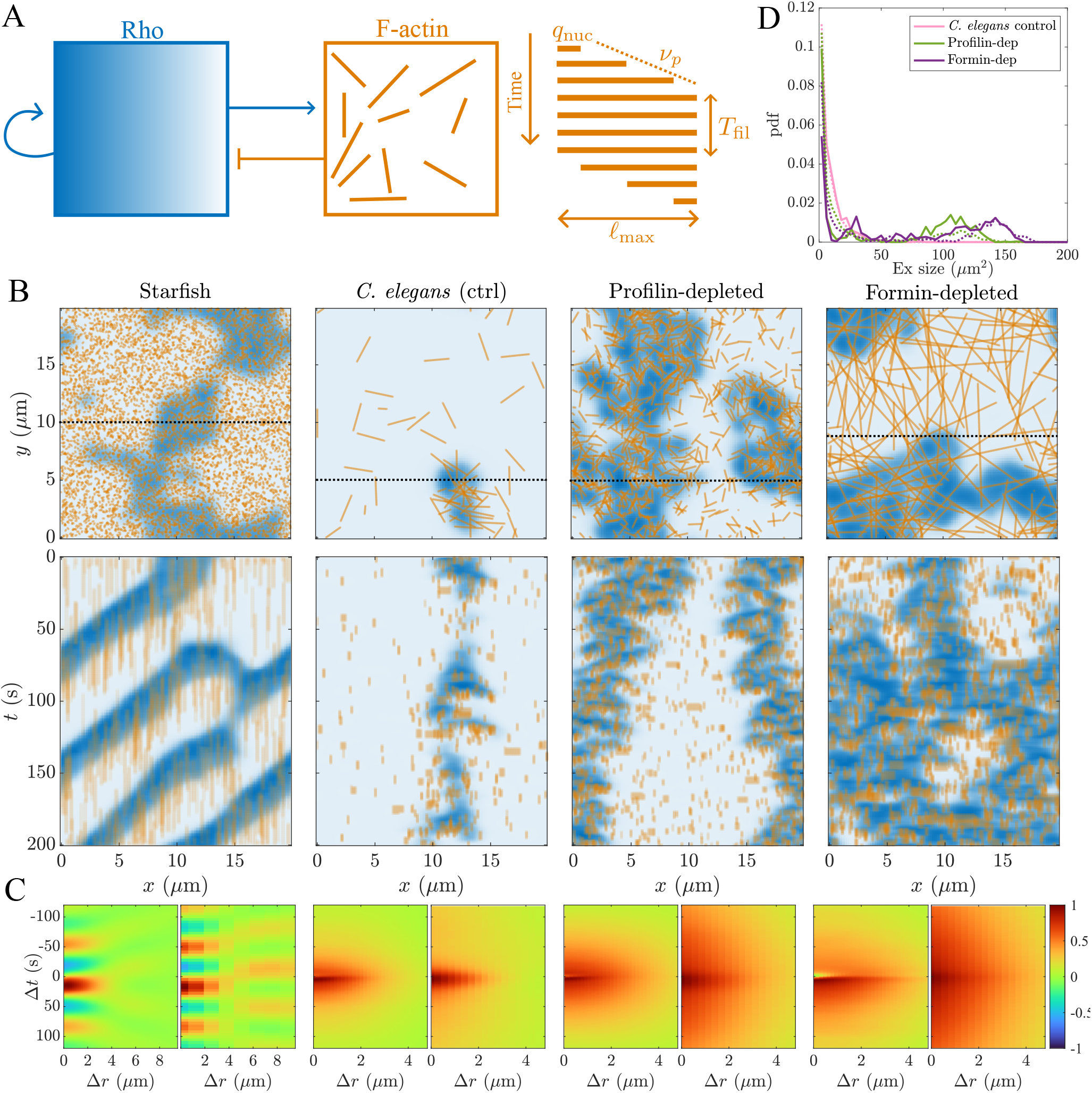
Hybrid discrete/continuum model reproduces experimental data across a variety of model systems and experimental conditions. (A) Model schematic. A continuum model of Rho is connected to a discrete actin network through activator-inhibitor coupling. Each discrete actin filament goes through the lifetime cycle shown at right: nucleation (rate given by (2b)), assembly (speed *ν*_*d*_ = *ν*_*p*_) to a fixed length (*𝓁*_max_), a fixed time (*T*_fil_) spent at that length, and disassembly from the older end (speed *ν*_*p*_). (B) Still images (at *t* = 0) and kymographs (along the indicated slices) for representative best-fit parameter sets for each experimental condition. (C) Cross correlations in the hybrid model (averaged over two simulations) compared to their experimental counterparts (right). (D) Excitation size distributions in the *C. elegans* parameter sets, compared to experimental data (dotted lines).

From the positions of the discrete filaments, we define a continuum F-actin field *f* (***x***, *t*) by evaluating the convolution with a Gaussian of width *g*_*w*_ on a grid (see (7) in Methods). We then assume enhanced inactivation of Rho by F-actin using the linear relationship

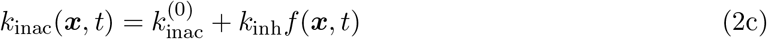

in (2a). The basal rate of Rho inactivation 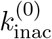 incorporates the combined effects of basal GAP activity and a homogeneous background actin network (e.g., Arp 2/3-mediated branched networks). The inhibition enhancement *k*_inh_*f* reflects the (heterogeneous) contribution of the explicit actin filaments.

### 2.2 Parameter estimation and inference

In the experimental systems that we examine, Rho dynamics are characterized by transitions between high and low activity states. To ensure that our model captures these dynamics, we choose the parameters in (2a) such that local Rho activity levels are close to a bifurcation (in the parameter *k*_inac_). As shown in Fig. S5, in the absence of diffusion, the local dynamics of Rho fall into three categories as a function of *k*_inac_. For low *k*_inac_, there is a single steady state with high Rho concentration. For high *k*_inac_, there is a single steady state with low Rho concentration. At intermediate *k*_inac_, the Rho concentration is bistable.

Based on the bifurcation diagram, there are two plausible choices for the local behavior of Rho in the absence of F-actin, which is set by the value of 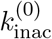 in (2c). For small 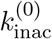, any void in the actin network automatically initiates a pulse of Rho. For intermediate 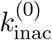, Rho is bistable in the absence of F-actin, and excitations are initiated only when Rho activity enters from neighboring regions (e.g., traveling waves). As we show in the SI (Section S2.4 and Fig. S7), the small 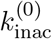 regime yields a negative cross-correlation between Rho and F-actin at (Δ*r*, Δ*t*) = (0, 0), as low actin concentration is always correlated with high Rho concentration. This correlation pattern is inconsistent with the experimental data (Fig. 1), which leads us to choose an intermediate 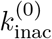 where excitations spread by diffusion (with diffusivity *D*_*ρ*_ = 0.1 *µ*m^2^/s taken from experimental measurements [74]). Since only the product *k*_inh_*f* appears in (2a), we also fix *k*_inh_ to make *f* unique. Finally, an empirical analysis (Fig. S8) shows that summary statistics are not sensitive to the individual polymerization and depolymerization rates *ν*_*p*_ and *ν*_*d*_, provided that the total time a filament spends growing and shrinking is conserved. For this reason, we set *ν*_*d*_ = *ν*_*p*_, reducing the unknowns to five F-actin assembly parameters: *q*_*b*_, *q*_*ρ*_, *ν*_*p*_, *T*_fil_, and *𝓁*_max_ (Fig. 2A), which we vary to explore how emergent behaviors depend on local F-actin assembly dynamics.

To estimate the parameters from data, we define a posterior distribution Π(***p***|***d***) that gives the probability of observing the parameters ***p*** for each data set ***d***. This distribution can be sampled by choosing parameter set ***p***, running simulations to obtain simulated data, and computing the error in cross correlation (for all conditions) and excitation sizes (for *C. elegans* embryos only). See Methods for a more formal description of our inference procedure.

### 2.3 A discrete/continuum model reproduces Rho dynamics across model systems

We first ask whether the model can reproduce the variations in experimental data by tuning F-actin assembly dynamics. To this end, we obtain 5 × 10^5^ independent samples of the model (simply by sampling uniformly within a bounded space, see Table S1), extract the parameter set with maximum posterior density for each data set, and visualize the model output in Fig. 2B.

Snapshots and kymographs reveal a strong resemblance to the experimental data (Fig. 1B). For the starfish data, the best fit simulations reproduce classical traveling waves, and the actin filaments are homogeneously distributed, with small lengths rendering their orientations superfluous. By contrast, best-fit simulations for *C. elegans* control embryos show actin filaments with lengths of a few microns, which create substantial heterogeneities in inhibitor dynamics. Snapshots and kymographs show how excitations escape through voids in the actin network, setting up a steady state of small, meandering patches of excitation which extinguish stochastically. In simulations that best match profilin- and formin-depleted embryos, these excitations grow in size, and cross correlations become broader in space and time. For simulations that best match profilin-depleted embryos, this occurs because filaments are shorter, which confines the spread of inhibition, allowing excitations to spread until they run into filaments nucleated independently of Rho. For simulations that best match formin-depleted embryos, filaments are long and assemble rapidly from excitations. These long filaments give the most heterogeneous architecture, with large gaps through which excitations spread.

### 2.4 Transforming inferred parameters into observables for model validation

To validate the model, we compare the inferred parameter values to available experimental measurements. Agreement between inferred and measured values suggests that the model has captured essential features of the relationship between F-actin assembly and emergent Rho dynamics. Otherwise, the inference problem may be ill-posed and more information may be required to constrain the parameters.

We transform the five F-actin assembly parameters into four observables that have been measured experimentally or could be in the future. The first observable is the turnover time of a monomer or subunit within a filament:

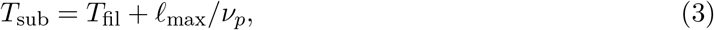

which can be measured by single-particle tracking of actin monomers bound to the cortex [37, 49, 74]. The second observable is the length of an actin filament *𝓁*_max_, which can be measured by tracking single formin molecules as they polymerize F-actin at the barbed end (and multiplying the polymerization speed by the lifetime) [72]. The filament length *𝓁*_max_ quantifies the lengthscale of inhibition, which competes with the Rho diffusion lengthscale 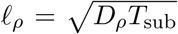 over the lifetime of an actin subunit.

The two other observables convert the basal and Rho-mediated nucleation rates into a density (length of filamentous actin per square length of domain) of basal and Rho-mediated actin filaments on the cortex. Using an effective filament turnover rate

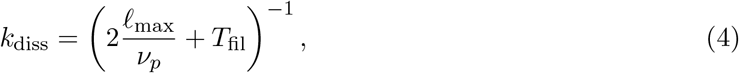

the total number density of basal and Rho-mediated filaments is given roughly by *N*_*b*_ = *q*_*b*_*/k*_diss_ and 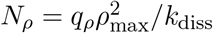, where *ρ*_max_ is the largest steady-state Rho concentration. In turn, the total length of basal/Rho-mediated actin per domain area is *L*_*b/ρ*_ = *𝓁*_max_*N*_*b/ρ*_. These quantities could be measured experimentally by computing the actin filament density in areas of high and low Rho activity, then solving a 2 × 2 system of equations to obtain *L*_*b*_ and *L*_*ρ*_.

Figure 3 shows the observables for the fifty parameter sets with maximum posterior density in each experimental condition. Comparing likely parameters for starfish and control *C. elegans* embryos (blue and pink symbols) reveals a sharp difference in the rate of actin filament turnover and heterogeneity. The curve 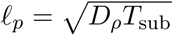 (dotted black) demarcates the boundary above which the spread of inhibition exceeds the spread of excitation. Parameters for starfish fall below this line, indicating that actin dynamics lag Rho dynamics, while parameters for *C. elegans* fall above it, indicating that actin dynamics control the spread of Rho. For starfish parameters, subunit turnover times cluster around 30 s, while for control *C. elegans* the lifetimes are about 5–15 s (left panel of Fig. 3). Remarkably, the inferred mean turnover times agree with independent experimental measurements of 40 s [75, Fig. 5F] and 9 s [37, 74]. The inferred filament lengths of a few microns are also similar to those measured in control *C. elegans* embryos (up to 6 *µ*m [72]). Thus, our model reveals that the excitation dynamics (specifically, the summary statistics of cross correlation and excitation size) in control *C. elegans* embryos actually *imply* the microscopic F-actin dynamics (at least, the ones that have been measured to date). This correspondence lends more confidence to the model’s prediction that faster actin turnover and longer filaments are responsible for the transition from traveling waves in starfish to transient pulses in *C. elegans*.

**Figure 3:**
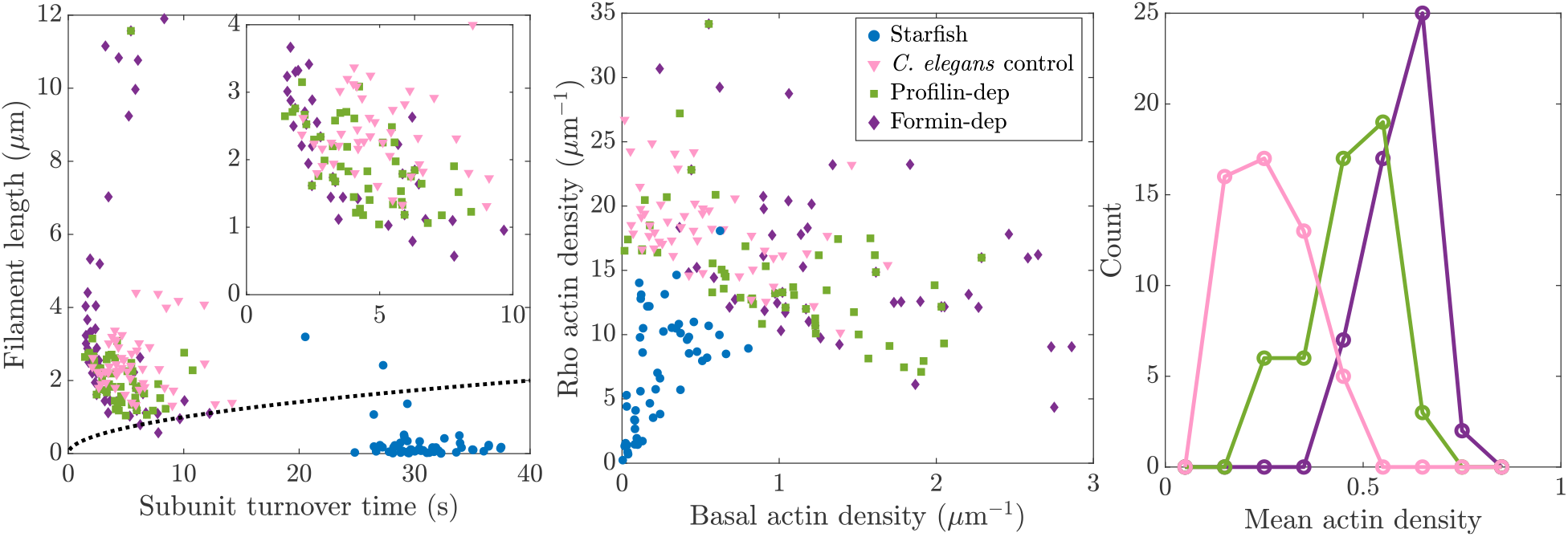
Hybrid model (constrained on cross correlations and excitation sizes) successfully infers actin kinetics in starfish and control *C. elegans* embryos but misses key features of formin and profilin depletion. We show a series of statistics for the 50 (out of 5 × 10^5^) highest-likelihood parameter sets in starfish oocytes (blue) and control (pink), profilin-depleted (green), and formin-depleted (purple) *C. elegans* embryos. Left panel: The subunit turnover time 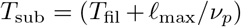 versus filament length *𝓁*_max_. The dotted line shows the intrinsic Rho lengthscale 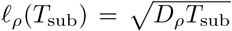, and the inset shows an expanded view of the bottom left corner. Middle panel: The density of actin from basal versus Rho-mediated nucleation. Right panel: Distribution of mean actin densities (total filament length per *µ*m^2^) across the fifty most likely simulations.

Previous work [37, 55, 70] showed that embryos depleted of profilin have longer turnover times and shorter filament lengths, while embryos depleted of either profilin or formin have significantly less overall F-actin. Our parameter inference matches some of these observations but not others; parameter sets for profilin depletion typically have shorter filament lengths (see green squares in Fig. 3), but these parameter sets also display shorter subunit turnover times than controls. Significantly, both the profilin- and formin-depleted embryo simulations have higher F-actin densities than controls (right panel of Fig. 3), in disagreement with experimental observations. Thus, in these cases, more information is required to constrain F-actin assembly parameters prior to interpreting model output.

### 2.5 Correcting F-actin dynamics with prior knowledge

The failure of the model to reproduce experimentally-relevant F-actin densities means that those observations are not embedded in the cross correlation and excitation sizes (which is the only information the model has). In fact, because the cross correlation between Rho and F-actin is positive everywhere in *C. elegans* embryos, data sets with the most Rho activity (formin-depletion) have the highest inferred F-actin concentration. Thus, we must incorporate what we already know about actin dynamics, which we do by modifying our inference procedure to include a penalty on unphysical parameter sets (see (6e) in Methods). In particular, we bias parameter sets to increase the amount of F-actin in simulations of control *C. elegans* embryos and decrease the amount of F-actin in simulations of formin- and profilin- depleted embryos. In simulations of profilin-depleted embryos, we also select for parameter sets with subunit turnover times *T*_sub_ that are roughly double those in controls. Table 1 summarizes the experimental observations included in the prior, as well as those already inferred from cell-scale data.

**Table 1:** Summary of known experimental observations on actin kinetics, and whether they are inferred by the model. If not, these observations are included in parameter constraints (prior knowledge).

| Experimental observation | Value | Inferred? |
| --- | --- | --- |
| Subunit lifetime, starfish | 40 s [75] | Yes |
| Subunit lifetime, *C. elegans* controls | 8 s [37] | Yes |
| Subunit lifetime, profilin-depletion | 16–32 s (unpublished) | No |
| Filament length, control *C. elegans* | Up to 6 $\mu\text{m}$ [72] | Yes |
| Actin densities, profilin/formin-depletion *C. elegans* | Reduced [55] | No |

Repeating our search through parameter space with the constrained F-actin dynamics, the bestfit parameter sets in control embryos now have more F-actin than profilin- and formin-depleted ones (Fig. 4A). Examining the best-fit simulations once again, control embryos have more actin filaments in regions of low Rho activity, which prevents much spread of excitations relative to the other conditions. The profilin-depleted phenotype is roughly unchanged, as filaments are significantly shorter than controls and excitations are larger (they spread instead of shrink). Penalizing the amount of F-actin completely removes the long filament phenotype previously observed for formin depletion; instead, we obtain excitations which readily grow and move through the regions lacking F-actin. As might be expected, the summary statistics (cross correlations and excitation sizes) do not line up as well in the simulations constrained on F-actin density. They do, however, reproduce the general trends of the data: excitation sizes are larger in formin and profilin depleted embryos, and cross correlations are wider in space (the spread in time appears roughly the same across all simulations).

**Figure 4:**
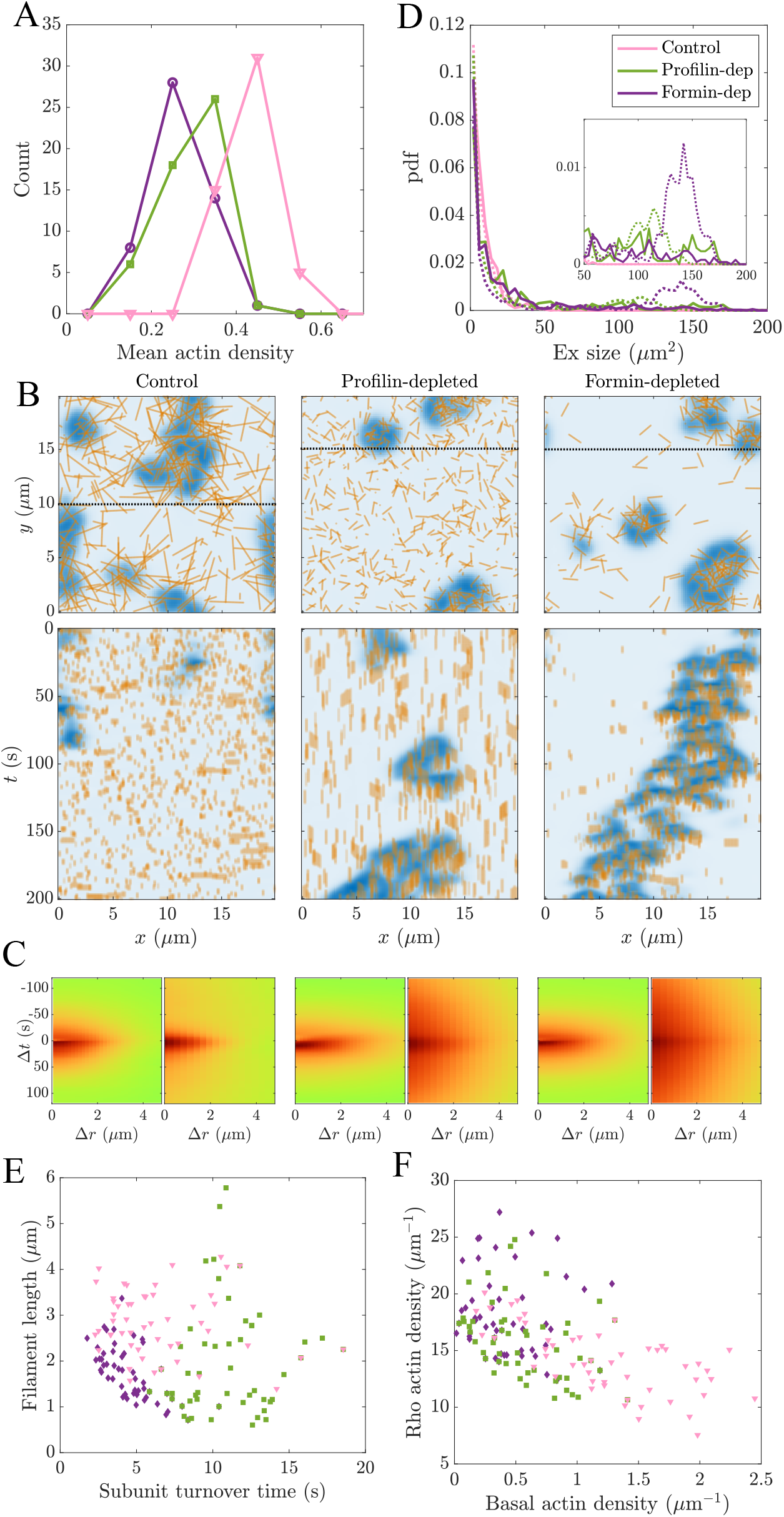
Constraining actin density and kinetics yields best-fit simulations that align with experimental observations. (A) F-actin densities for the fifty best-fit parameter sets obtained when enforcing the ordering of mean F-actin density (see (6e)). (B) Most likely parameter set for each data set (starfish results are unchanged from Fig. 2 and omitted). Top plot shows snapshot at *t* = 0, bottom plot shows kymographs over the indicated slices. (C) Cross correlations and (D) excitation size distributions for the simulations shown in (B); inset magnifies sizes in range [50, 200]. (E,F) Fifty most likely parameter sets (highest posterior densities), using the same observables as Fig. 3.

In cases where the unconstrained model produced unrealistic results for F-actin density, mechanistic insights can only be obtained when model parameters are constrained. Examining the fifty constrained parameter sets with highest posterior density (Fig. 4E,F), control embryos retain the characteristic turnover rate of 5–15 s. The long-filament phenotype is removed from the formin-depleted parameter sets (because the F-actin density is too high), and filaments are shorter with a slightly faster turnover rate than controls (Fig. 4E). Profilin-depleted parameter sets typically have shorter filament lengths than controls as well (green squares in Fig. 4E).

Perhaps the most interesting effect of constraining the actin dynamics is the clear trend that emerges in the basal versus Rho-mediated F-actin density plot (Fig. 4F). Control embryos have high basal F-actin densities and low Rho-mediated F-actin densities, while profilin- and formin-depleted embryos have low basal F-actin densities and high Rho-mediated actin densities. This comparison suggests that reducing the amount of basal actin and increasing the amount of Rho-mediated F-actin favors larger and more stable excitations.

## 3 Extracting the minimal mechanisms behind spatiotemporal variation in Rho dynamics

Our hybrid model reproduces the variation in Rho dynamics observed across experimental systems, but the additional microscopic complexity makes it difficult to determine the reason(s) for this success. To address this issue, we seek to lift key features of the discrete filament dynamics into a continuum model and determine how each feature shapes the predicted Rho dynamics. We identify two features (Fig. 5A) of the discrete F-actin dynamics whose contributions are not captured by classical reaction-diffusion models [14]. First, the dynamics of barbed-end assembly and pointed-end disassembly results in effective directional transport of F-actin density and orientation, which manifests as advective, rather then diffusive, transport in a continuum-level description. Second, small numbers of filaments in local regions of the cortex imply stochastic variation in the rates and orientations of F-actin assembly, and thus in the magnitude and direction of advective transport.

Since oriented F-actin assembly is key to both transport and stochasticity, we introduce a continuum formulation where the F-actin density *f* (***x***, *θ*) is a function of both position and orientation [62, 76–80]. We approximate the discrete growth and shrinkage dynamics as advection along ***τ*** (*θ*) = (cos *θ*, sin *θ*), with speed taken from the hybrid model parameters as *ν*_*p*_ = *𝓁*_max_*/T*_sub_. Combining this with local assembly (i.e., nucleation and growth) and disassembly gives the evolution equation

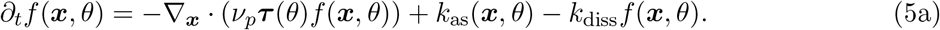

The dynamics of Rho are still governed by (2a) and (2c), but with *f* replaced by the total amount of F-actin at each point in space,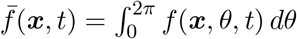.

### 3.1 Isotropic advective transport mimics classical diffusion, failing to explain variations in Rho dynamics

We first ask if advective transport alone, without any stochasticity, can reproduce the full range of Rho dynamics. In this case, F-actin assembly occurs isotropically with rate

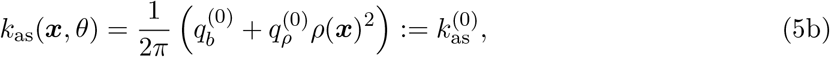

chosen to mimic the hybrid model. Simulating the dynamics (5a) and (2a) with turnover (1*/k*_diss_ = 30 s) and transport (*ν*_*p*_ = 0.025 *µ*m/s) rates consistent with hybrid model fits to the starfish data (see Table S3) yields traveling waves with an oscillatory cross correlation (Fig. 5B and B’), reproducing the experimental dynamics. However, when the transport (*ν*_*p*_ = 0.5 *µ*m/s) and turnover rates (1*/k*_diss_ = 8 s) are increased to match the hybrid model parameters for *C. elegans* controls, standing patterns of Rho excitation result, in disagreement with experimental behavior.

The failure of the modified continuum model in (5a) to reproduce the transient mobile excitations of Rho observed in *C. elegans* reflects the qualitative similarity between isotropic advective transport and classical diffusion. In the model in (5a), the preferred orientation of F-actin (shown using arrows in Fig. 5) points radially outward from Rho excitations (Fig. 5C). Because F-actin assembly rates increase with local Rho activity, isotropic assembly and advection implies a flux of F-actin down gradients of Rho activity, analogous to classical diffusion. Fast F-actin transport down Rho gradients “cages in” excitations, while slow F-actin transport trails spreading waves of Rho activity. Therefore, a continuum model with isotropic advection of F-actin produces the same range of behaviors as a continuum model with isotropic diffusion (Fig. S9), which fails to include transient excitations characteristic of *C. elegans*.

**Figure 5:**
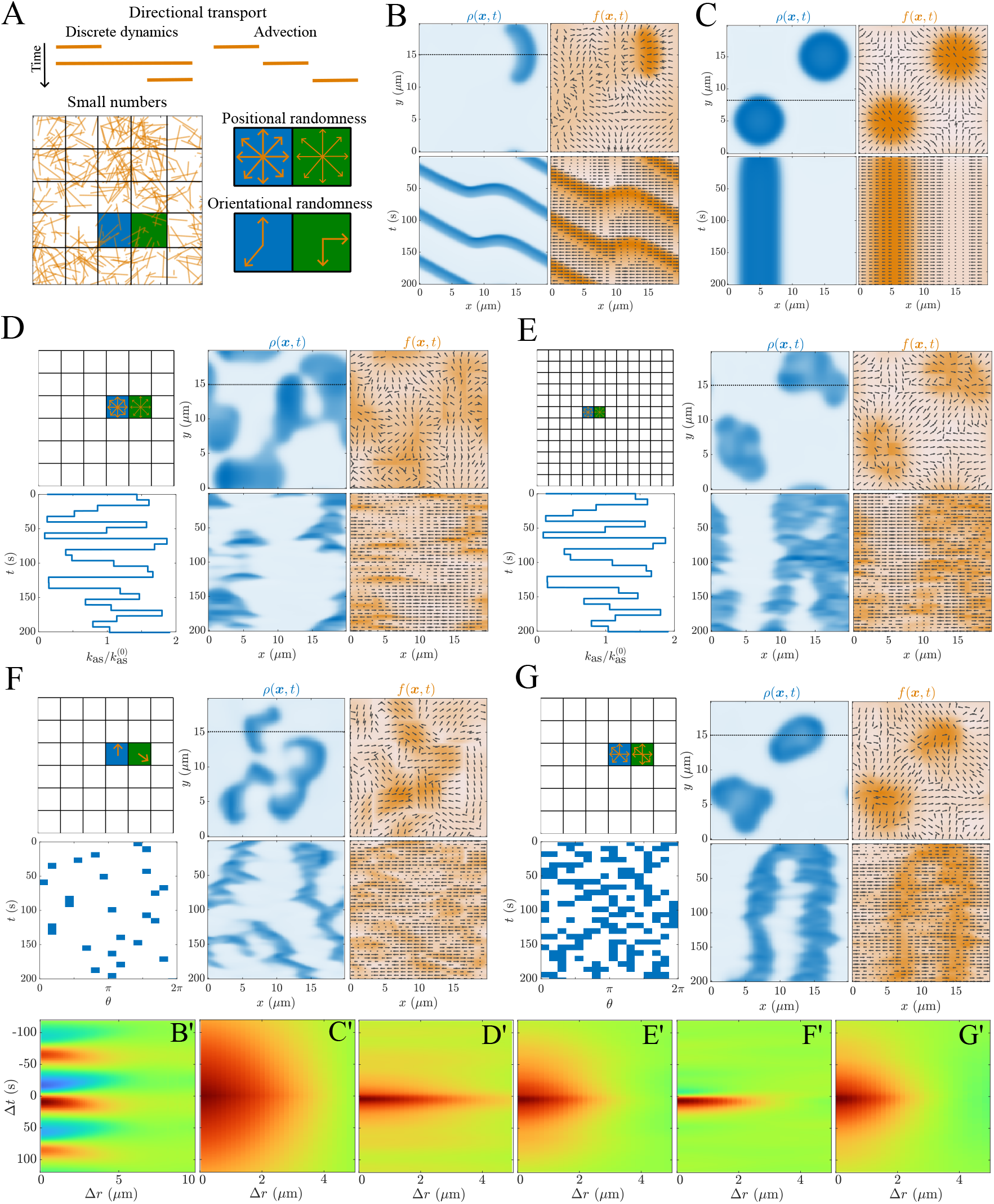
Adding spatial and orientational stochasticity enables a continuum model to reproduce both traveling waves and transient pulses. (A) Extracting the essential features of the hybrid model dynamics in a continuum model. The discrete polymerization dynamics are similar to advective transport, and small numbers imply local stochasticity in orientation and number within local regions of the cortex. (B–G) Simulations and (B’–G’) cross correlations of the continuum F-actin and Rho dynamics, (2a) and (5a). (B–C) Without stochasticity (constant assembly rates given by (5b)). (B) Slow F-actin turnover (1*/k*_diss_ = 30 s) and transport (*ν*_*p*_ = 0.025 *µ*m/s). (C) Fast F-actin turnover (1*/k*_diss_ = 8 s) and transport (*ν*_*p*_ = 0.5 *µ*m/s). (D–E) With positional stochasticity on regions of the domain (assembly rates given by (5c)). Plots at top left show regions; plots at bottom left show the normalized nucleation rates on a sample region (the others vary similarly and independently). (D) Region side length 3.3 *µ*m. (E) Region side length 1.7 *µ*m. (F–G) With orientational stochasticity (assembly rates given by (5d)). Top left plots show the regions, bottom left plots show the angle(s) nucleated on each region over time. (F) One angle nucleated on each region. (G) Six angles nucleated on each region. In the actin snapshots, arrows indicate the most likely orientation of F-actin ***τ*** (*θ*_max_), with the *x* projection of this orientation shown in kymographs.

### 3.2 Full range of Rho dynamics can be reproduced when spatial and orientational randomness create heterogeneities in F-actin inhibition

We next ask whether stochastic fluctuations in local rates or orientations of filament assembly, which are embedded in the hybrid model, are key to producing transient mobile patterns of Rho excitation. To capture these stochastic effects in a continuum model, we divide the domain into square regions ***A***_*i*_ which have mean size characteristic of individual actin filaments, on which we distinguish two types of randomness (Fig. 5A). First, we consider *positional* randomness, where the rates of assembly are isotropic but vary randomly across regions (Fig. 5(D–E)). Then, we consider *orientational* randomness, where the rates of assembly are anisotropic but constant in magnitude across regions (Fig. 5(F–G)).

#### 3.2.1 Large-scale spatial heterogeneity makes excitations transient

To implement positional randomness, we scale the (isotropic) assembly rates by a region- and time-dependent constant, so that

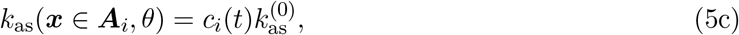

for *c*_*i*_ chosen randomly and uniformly on [0, 2], and resampled every 1*/k*_diss_ to capture the timescale of filament turnover (Fig. 5D shows 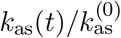 in one of the regions). Using regions with side length 3.3 *µ*m (corresponding to the scale of the inferred filament length in the hybrid model), and *k*_diss_ = 8 s to reflect fast F-actin turnover measured in *C. elegans*, the excitations become transient (Fig. 5D), and the cross correlations sharpen, becoming similar to control *C. elegans* embryos (Fig. 5D’). The appearance and disappearance of Rho pulses are the consequence of stochastic effects [56], in which local increases in F-actin assembly destroy excitations, while local decreases create gaps into which excitations can spread. Still, because transport of F-actin is fast and isotropic, the spread of excitation is relatively limited (kymograph in Fig. 5D).

Under the same conditions, halving the lengthscale of spatial randomness gives larger, more persistent patches, and a broader cross correlation (Fig. 5E). Indeed, the randomness averages out on the lengthscale of Rho excitations, resulting in patterns that are more similar to the isotropic advective case (Fig. 5C). This explains why shorter filament lengths were inferred in formin- and profilin-depleted embryos, where excitations are larger and more persistent.

#### 3.2.2 Significant orientational heterogeneity makes excitations mobile

To implement orientational randomness, we choose a random sample **Θ**_*i*_ of *n*_*θ*_ possible orientations to assemble on each region, then set

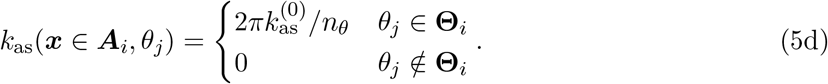

To capture the timescale of filament turnover (thus orientation persistence), we resample **Θ**_*i*_ on each region every 1*/k*_diss_ = 8 s (Figure 5(F,G) shows a heatmap of the discrete angles assembled in one region over time).

When only one orientation is assembled on each region, standing patterns once again break up into transient excitations, but this time with significant mobility (Fig. 5F). Transport of Rho into regions that lack F-actin is evident from the more negative cross correlation on (Δ*r*, Δ*t* < 0) (Fig. 5F’), and from kymographs (Fig. 5E) showing no spread of Rho in the direction of the preferred Factin orientation (arrows). Indeed, when only a subset of F-actin orientations is assembled locally, the rest of the directions are available for Rho diffusion.

In profilin- and formin-depleted embryos, the hybrid model inferred more Rho-mediated nucleation (relative to basal nucleation) and shorter filament lengths, which together imply an increase in the number of assembled orientations in regions of Rho excitation. Similar to the case of spatial randomness, we find that accounting for more homogeneous F-actin assembly in the continuum model (by assembling six angles per region instead of one) transitions the dynamics back toward standing patterns with broader cross correlations (Fig. 5G and G’). Assembling more orientations in regions of Rho excitation gives more directions of filament growth, but also less growth in a particular direction (since the total nucleation rate is fixed). This simultaneously limits the spread and break up of excitations, leading to larger and more stable patterns.

#### 3.2.3 Combining spatial and orientational heterogeneities reproduces the full range of dynamics

True F-actin assembly dynamics, both *in vivo* and in the hybrid model, induce both spatial and orientational heterogeneity. Including both forms of randomness simultaneously in the continuum model can reproduce the full range of observed behaviors (Fig. S10), from starfish (where randomness has the least impact) to control *C. elegans* embryos (where randomness has the most impact) to profilin- and formin-depleted *C. elegans* embryos (intermediate impact).

## 4 Discussion

How functional diversity in cellular behaviors could arise through different “tunings” of conserved systems of molecular interactions remains a major question in biology. Here, we focused on a conserved system of activator-inhibitor interactions between active Rho and actin filaments at the cell cortex. How these components produce traveling waves in starfish and frogs and transient pulses in worms has not been clear from reaction-diffusion models. Based on previous experiments [55,75], we hypothesized that the differences between the two systems could come from the underlying F-actin dynamics. Introducing a hybrid model that coupled discrete actin filament dynamics to continuum dynamics of Rho, we demonstrated how changes in F-actin heterogeneity and stochasticity lead to the observed changes in Rho activity.

While the experimental data [21, 55] and modeling techniques [14, 49, 61, 79–82] all existed previously, the novelty of our approach comes from the close interaction between the experiment and modeling. By extracting a set of summary statistics (cross correlations and excitation sizes) from the data, we formulated a posterior probability distribution, which we maximized to infer the parameter sets responsible for each phenotype. Without any prior knowledge, the model initially predicted rates of F-actin turnover and filament lengthscales that agreed with previous measurements in starfish and *C. elegans* controls [37, 74, 75]. When fitting the data for formin- and profilin-depleted embryos, which have larger excitations and diffuse cross correlation profiles, the model initially inferred parameters inconsistent with expectations. Thus, in these embryos, cross correlation and excitation size alone do not supply sufficient information to infer the correct F-actin dynamics. To address this issue, we introduced a constraint on the amount of F-actin, under which the model inferred shorter filaments and less basal actin assembly in profilin- and formin-depleted embryos. This result provides a prediction for further model validation, which can be tested experimentally by obtaining more detailed measurements of filament length and density, in concert with reducing measurement noise in the data (Fig. 1(C–E)).

The combination of hybrid and continuum modeling provided a mechanistic picture of how F-actin assembly dynamics shape the spread of Rho activity *in vivo*. Rho acts as a spreading wavefront that is confined by basal F-actin (F-actin assembled independently of Rho) and chased by elongation of Rho-mediated filaments (Fig. 6). Starfish have homogeneous distributions of slowly elongating filaments, which assemble behind traveling wavefronts of Rho excitation. *C. elegans* embryos, by contrast, have actin filaments which elongate more rapidly, making the spread of inhibition faster than the spread of excitation. In classical continuum models of the Rho/F-actin circuit, faster spread of inhibition can only produce standing patterns, as actin filaments rapidly polymerize outward from excitations to cage them. However, our models revealed that randomness can create gaps and heterogeneities in the actin network, which cause excitations to become transient (spatial randomness) and traveling (orientational randomness), as observed in *C. elegans*. These excitations spread until Rho-mediated F-actin catches up with them, or until they run into actin filaments nucleated independently of Rho. These are not traditional traveling waves; because the rapid assembly of F-actin still dominates in most directions, a positive cross correlation between Rho and F-actin results at all times. While the same mechanism operates in formin- and profilin-depleted embryos, shorter filament lengths and more Rho-mediated nucleation (relative to basal nucleation) make the F-actin dynamics under these conditions more homogeneous, which results in larger, more stable excitations and broader cross correlations. Reduced basal actin nucleation also allows excitations to grow larger and spread faster, potentially making the dynamics more wave-like, as was observed previously [55].

**Figure 6:**
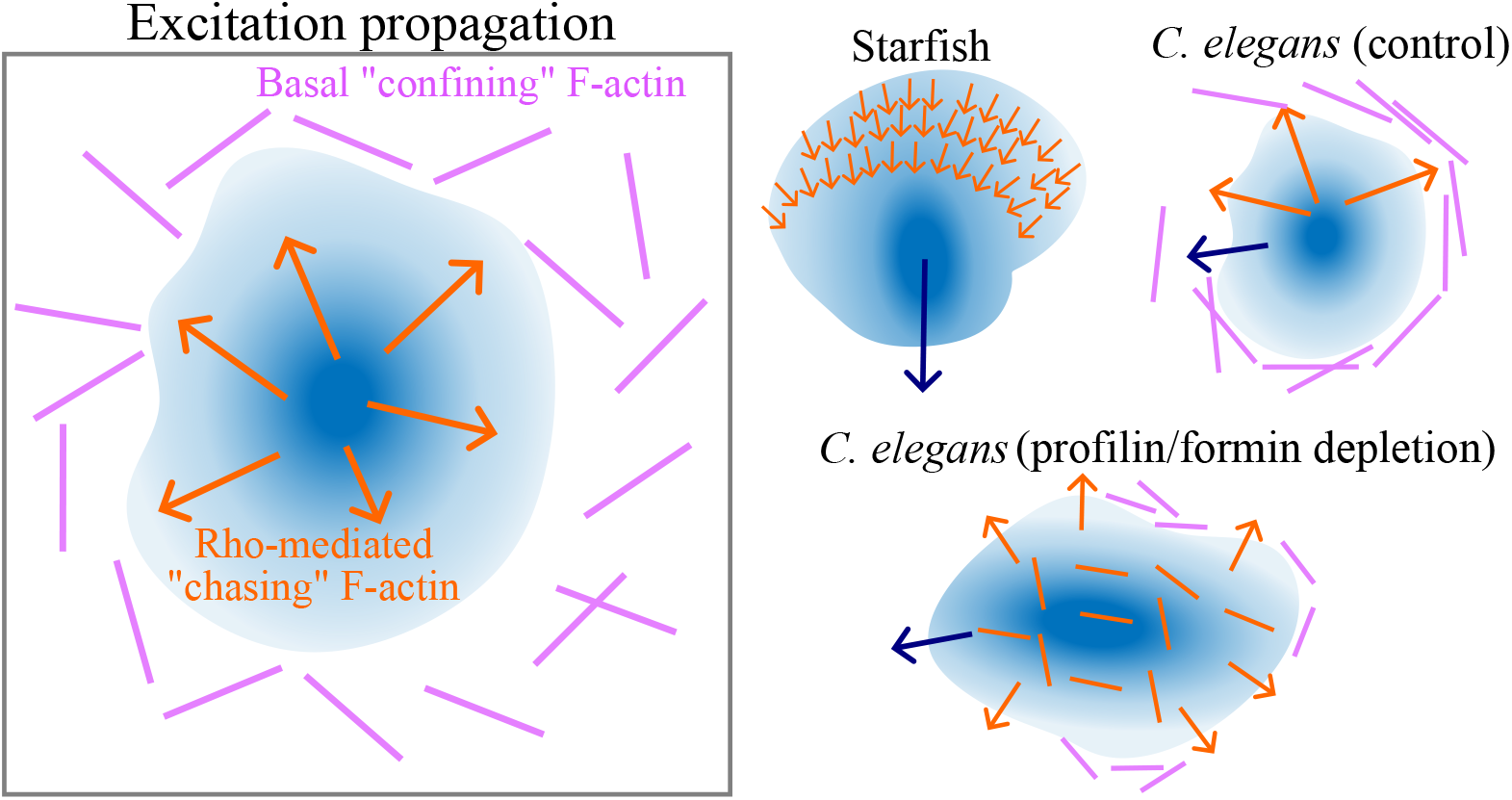
Schematic diagram demonstrating how basal and Rho-mediated F-actin confine and chase excitations, and how this process is adapted in starfish and *C. elegans* to produce pattern variation. In all cases, the active Rho wavefront spreads by diffusion, confined by “basal” F-actin (nucleated independent of Rho, purple) and chased by Rho-mediated assembly of F-actin (orange). In starfish embryos, low basal activity and slow F-actin elongation lead to traveling waves of Rho activity (navy arrow shows direction of Rho spread). *C. elegans* embryos have significantly higher rates of F-actin assembly, but more heterogeneous F-actin; this limits the size of excitations while allowing spread through (small) network voids. In profilin- and formin-depleted embryos, filaments are shorter and more homogeneous, leading to larger and more stable excitations, which spread through voids created by reduced basal assembly.

This work establishes our ability to infer parameters of microscopic actin assembly dynamics from cell-scale concentration data. This opens up an exciting future direction: using Rho GTPase and F-actin concentrations as probes to learn how various actin-binding proteins alter F-actin dynamics. By analyzing depleted versus control embryos, our model could predict how annilin [73], septin [66], or anterior PARs [83, 84] affect F-actin assembly dynamics through analysis of Rho GTPase excitation patterns and F-actin concentration fields. More generally, our approach could be used to infer how individual molecular players shape fundamental processes, such as cell division and migration, by modulating cytoskeletal dynamics [85, 86]. Accounting for the full spectrum of behaviors requires incorporating myosin-mediated contractility [55, 87, 88] into our discrete [89, 90] and continuum [39,50] models. The interplay between biochemistry and contractility will open up a broader range of possible excitation patterns and principles for self organization [39,40,51,52,91,92].

Our work also leads to new directions in mathematical analysis and computational science that go beyond the actin cytoskeleton. By using our discrete simulations to re-inform a continuum model, we demonstrated the importance of inhibitor heterogeneities—specifically spatial and orientational noise—in shaping the overall dynamics of activator-inhibitor systems. At this continuum level, a first-principles set of equations for anisotropic inhibition (obtained via homogenization theory [93, 94]) would significantly advance our understanding of self-organization in many biological systems. Applying our methodology to larger and more complicated biological systems will require methods that can efficiently sample high-dimensional parameter spaces. One outstanding challenge is to formulate a robust likelihood function for chaotic systems, optimizing summary statistics for chaotic trajectories [29, 95–98], while a separate challenge is to obtain sufficient sampling. Methods that compute an “amortized” or approximate likelihood (potentially via machine learning) seem particularly promising because they avoid repeated calls to a computationally-expensive forward model [28, 99–101]. Using these and other methods [102, 103] to leverage the large volumes of *in vivo* imaging data now available will enable us to deploy the approaches described here at scale.

## Data availability

Code for simulations and data analysis is available at https://github.com/omaxian/RhoActinRD

## Acknowledgments

All simulations were carried out on the Research Computing Center Midway cluster at the University of Chicago. OM was supported by the Yen and Chicago Fellows program, and is now supported by a Spark Fellowship from the Physics Frontier Center for Living Systems (CLS) funded by the National Science Foundation (PHY-2317138); ARD and EM also acknowledge support from the CLS.

## Methods

### Data analysis

For the starfish data, we select a region in the embryo interior and perform difference subtraction to remove static Rho signal [21]. For the *C. elegans* data, we consider *N* = 3 embryos for each condition, cut out a 20 *µ*m × 20 *µ*m section in the embryo interior, then correct for photobleaching by adjusting the mean intensity of every frame to be equal to the global mean. We filter the data in space and time using a Savitzky-Golay filter with window size 50 pixels/frames (5 *µ*m/30 s), and compute cross correlations on the filtered data (see Figs. S1–S2). To identify excitations in the filtered data, we set a threshold for excitations equal to 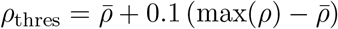, where 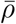 is the mean Rho level, and the maximum is over all (filtered) pixels in space and time. We then use standard image processing algorithms (bwconncomp in Matlab) to identify connected regions of excitation and make a histogram of excitation sizes (see Figs. S2–S3).

### Data-driven posterior estimation

Using Bayes’ theorem, the posterior **Π**(***p***|***d***) can be related to the likelihood **ℒ** (***d***|***p***) and prior **𝒫** (***p***) by

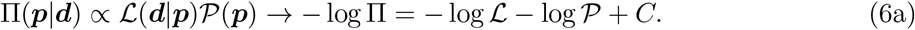

The prior assigns low probability to unphysical parameter sets, while the likelihood function quantifies the errors in the summary statistics of cross correlation and excitation size:

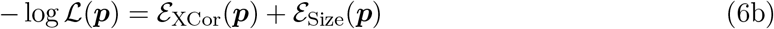

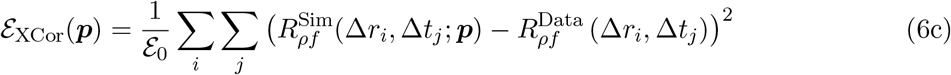

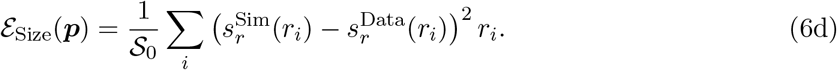

In (6c), 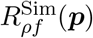 and 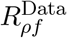 denote the simulated and experimental cross correlations, respectively, ℰ_0_ is set so that ℰ_XCor_ = 1 when 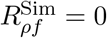, and the simulated cross correlation is resampled at the experimental data points by bilinear interpolation. Likewise, in (6d), 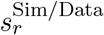 denotes the PDF of excitation sizes from simulations/data, and *S*_0_ is again set so that ℰ_Size_ = 1 when there are no excitations. The weight of *r*_*i*_ ensures that excitations are counted based on overall area and not absolute number; i.e., one excitation of size 200 *µ*m^2^ weighs the same as 200 excitations of size 1 *µ*m^2^. This gives a more faithful representation of errors in systems with large excitations (which are fewer in absolute number and therefore easier to discount without the weights). Finally, the *ℰ*_Size_ term is omitted for the starfish data, as the cross-correlation function in that case is sufficiently long-ranged in space and time to convey all of the necessary information.

### Prior densities

In initial parameter fitting (Fig. 2), we use a uniform prior on the parameter ranges given in Table S1. Subsequently (Fig. 4), we correct the posterior distribution using a prior on the actin density. Letting 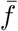 be the mean actin density, we penalize a lack of actin in control embryos and a surplus of actin in depleted embryos by setting

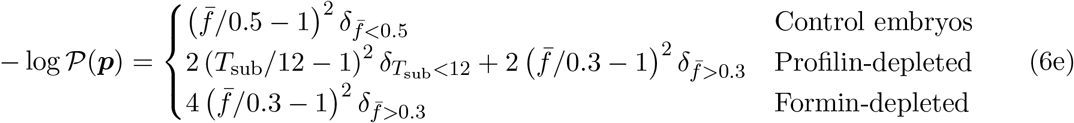

where *δ*_*x*_ is 1 if condition *x* is true and 0 otherwise. In addition to the density of actin, for profilin-depleted embryos we penalize subunit lifetimes shorter than 12 s, which is roughly double the average value for controls.

### Numerical methods

The continuum partial differential equations (2) and (5a) are solved numerically using a combination of a psuedo-spectral method (for diffusion) with an implicit-explicit temporal discretization, where diffusion is handled implicitly (using the backward Euler method), and the reaction terms are handled explicitly (using forward Euler). In the directional transport equation (5a), *θ* is discretized, and within a time step each direction is evolved separately, with advection handled explicitly via first-order upwinding. See [104, c. 1] for a description of these methods.

In the simulations with a discrete actin network, the actin dynamics occur in two steps: nucleation and polymerization. For nucleation, at each time step a filament nucleates in a grid cell centered at (*x*_*i*_, *y*_*j*_) if *r* < *q*_nuc_(*x*_*i*_, *y*_*j*_)Δ*x*^2^Δ*t*, where *r* ∼ *U* (0, 1). Once nucleated, the filament is assigned a random tangent vector (normalized Gaussian) and random (uniform) start point in the box (*x*_*i*_, *y*_*j*_)*±*(Δ*x*, Δ*y*). Polymerization and depolymerization thereafter are deterministic, where at each time step fibers grow/shrink by *ν*_*p*_Δ*t* length units, discretized into points ***X***_*j*_ with spacing Δ*s* (in the case when this results in a non-integer number of points added per time step, the remainder is handled stochastically such that the average growth/shrink rate is *ν*_*p*_Δ*t*). The associated actin filament field *f* (***x***) is defined by smearing the filaments with a Gaussian of width *g*_*w*_,

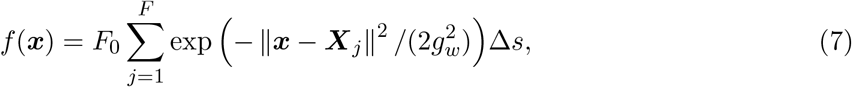

The constant *F*_0_ (units of concentration per length) is chosen so that the actin field on the grid is 1 for a single filament. We implement fast Gaussian gridding [105, Sec. 3] to accelerate the sum (7).

