## Supplemental Text and Figures for "Actin network heterogeneity tunes activator-inhibitor dynamics at the cell cortex"

### Supplemental Information for Actin network heterogeneity tunes activator-inhibitor dynamics at the cell cortex

**S4 Supplemental movie legends** **13**

#### S1 Data analysis

This section covers the conversion of raw experimental data from *C. elegans* into statistics of cross correlation and excitation size distribution. For each embryo, we focus analysis on a 200 pixel $\times$ 200 pixel (20 $\mu$m $\times$ 20 $\mu$m) region in the embryo interior. We first correct for photobleaching by adding/subtracting a frame-dependent constant intensity so that the mean of each frame matches the global mean. Then, we filter the input data before computing the cross correlation function. Finally, we perform thresholding to identify excitations.

##### S1.1 Filtering

We systematically examine the effect of filtering on data from control embryos (myosin(RNAi)) in Fig. S1. We consider two different filter types: a Fourier filter (which simply removes all high-frequency modes), and a Savitzky-Golay (SG) filter, which keeps some of the higher-frequency information. For larger numbers of modes and lower window sizes (20 Fourier modes and an SG window size of 20), the images look qualitatively similar. When the images are further smoothed (10 Fourier modes and an SG window size of 50), the Savitzky-Golay filter appears to capture a more faithful representation of the original data (the left panel is a good example of this). To smooth the noisy data while remaining faithful to the original excitation patterns, we choose a Savitzky-Golay filter with a window size of 50 in all dimensions (two spatial dimensions of 5 $\mu$m each and one time dimension of 30 s). The resulting smoothed data for each experimental condition are shown in Fig. 1 of the main text. These are used as inputs for the cross correlation function.

##### S1.2 Identifying excitations

To obtain discrete excitations from the filtered data, in each movie we identify a threshold equal to

$$\rho_{\text{thres}} = \bar{\rho} + \chi \left( \max(\rho) - \bar{\rho} \right), \tag{S1}$$

where $\bar{\rho}$ is the mean Rho level of that movie and the maximum is over all (filtered) pixels in space and time. Here $\chi$ is a dimensionless parameter which controls the fraction of the range (between the mean and maximum) at which we threshold. To choose it, we examine the identified excitations in the filtered data (Fig. S2). When the dimensionless threshold parameter $\chi$ is too low, the procedure picks up too many small excitations. When $\chi$ is too high, we miss too many of the excitations. We therefore choose an intermediate value, $\chi = 0.1$, to roughly match excitations to those observed by eye. After identifying regions of excitation, we use standard image processing algorithms (`bwconncomp` in Matlab) to separate different connected regions of excitation, then make a histogram of excitation sizes (Fig. S3).

##### S1.3 Comparing myosin-depleted to control embryos

To demonstrate that embryos depleted of myosin II manifest the essential features of control embryos, we compare summary statistics generated from *nmy-2*(RNAi) (used as controls in the main text) to *spd-5*(RNAi) embryos. As discussed in detail in previous work [49,55], *spd-5*(RNAi) is used to prevent the bulk cortical flows that occur during embryo polarization, thus allowing analysis of Rho and F-actin dynamics in a homogeneous background. Filtered snapshots and kymographs from these embryos strongly resemble *nmy-2(RNAi)*, and the summary statistics are indistinguishable between the two cases (Fig. S4).

#### S2 Hybrid model

In this section, we give further details on the hybrid model that couples a continuous Rho PDE with a discrete actin network.

##### S2.1 Parameters of Rho activation

The evolution of Rho is given by (2a) in the main text. For all of our simulations, we fix $K_{\text{sat}} = 0.1$, $k_{\text{b}} = 0.05$, and $k_{\text{ac}} = 1$. As discussed in Section 2.2 of the main text, in the absence of diffusion this

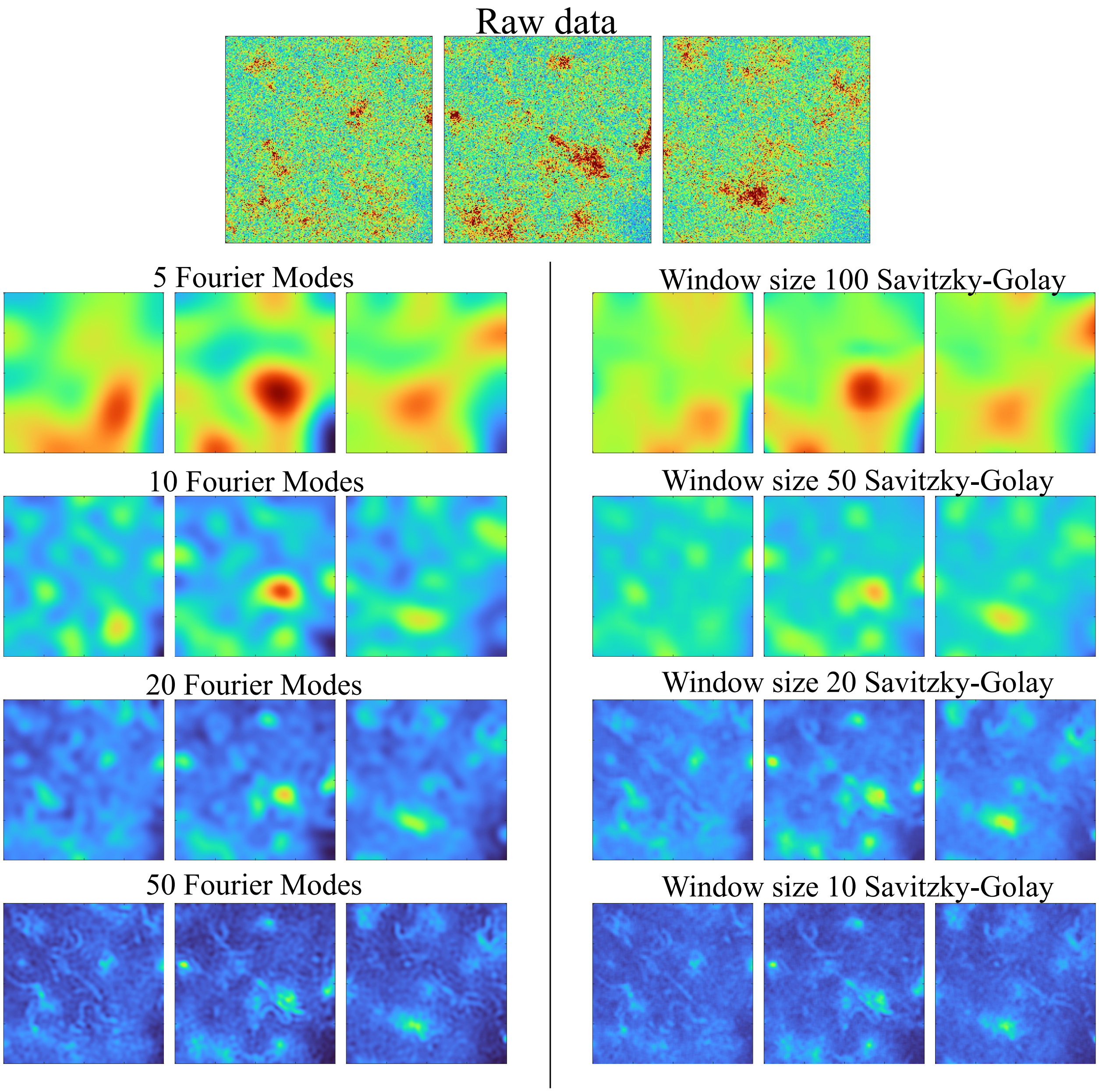

**Figure S1:** Filtering Rho dynamics in control (myosin(RNAi)) embryos. The top panels show the experimental data at three different times. The left column shows the resulting images filtered in all three dimensions (two spatial and one temporal) using Fourier filtering with the indicated number of modes remaining. The right column shows the filtered images using Savitzky-Golay filtering with the indicated window size. Each image is $20 \times 20$ $\mu$m.

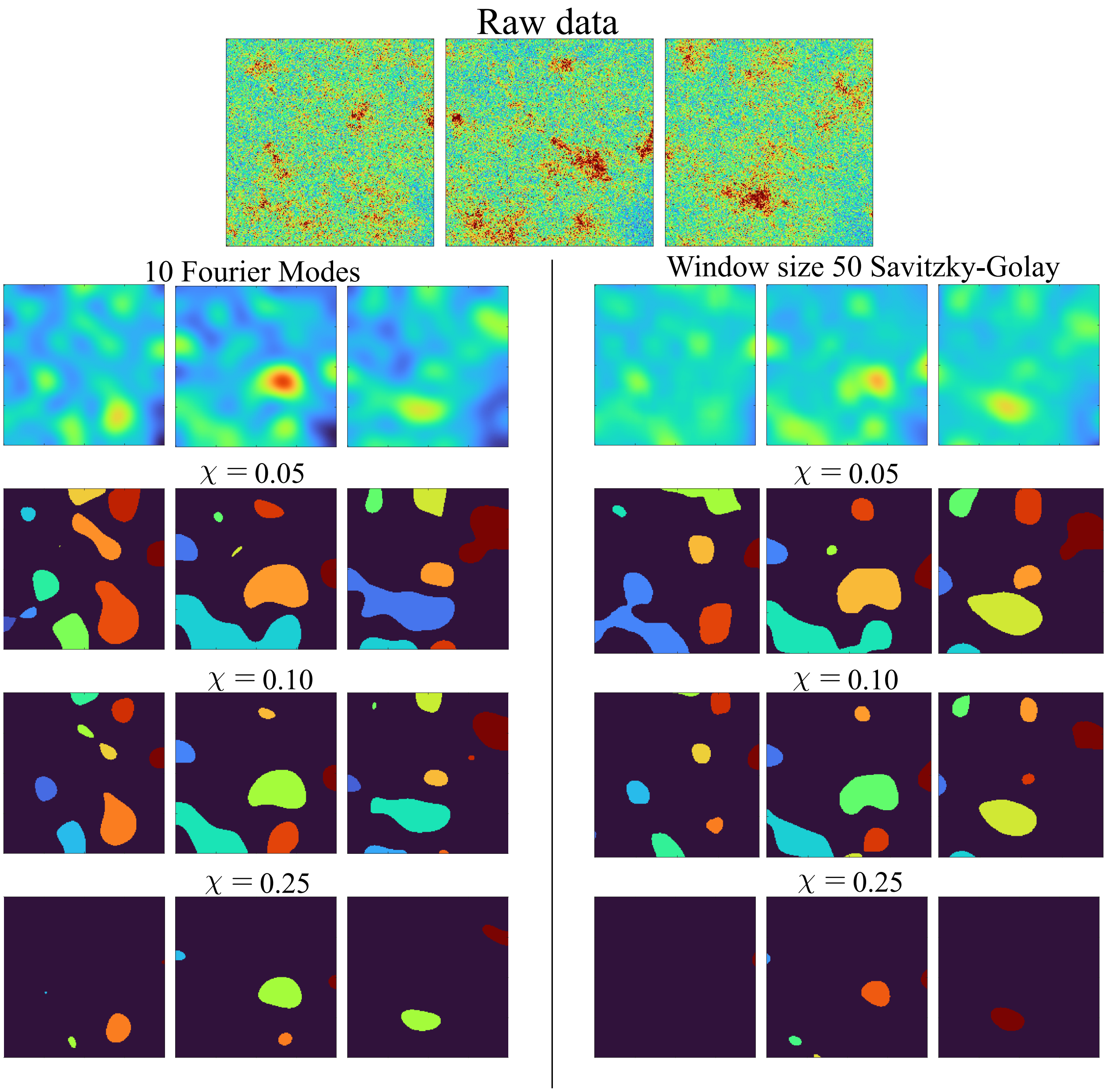

**Figure S2:** Identifying Rho excitations in control (myosin(RNAi)) embryos using filtering. The top panels show the experimental data, while the second row of (smoother) panels repeats the 10 Fourier mode / 50 Savitzky-Golay window size from Fig. S1. Subsequent panels show the identified excitation regions with various values of the threshold parameter $\chi$ in (S1).

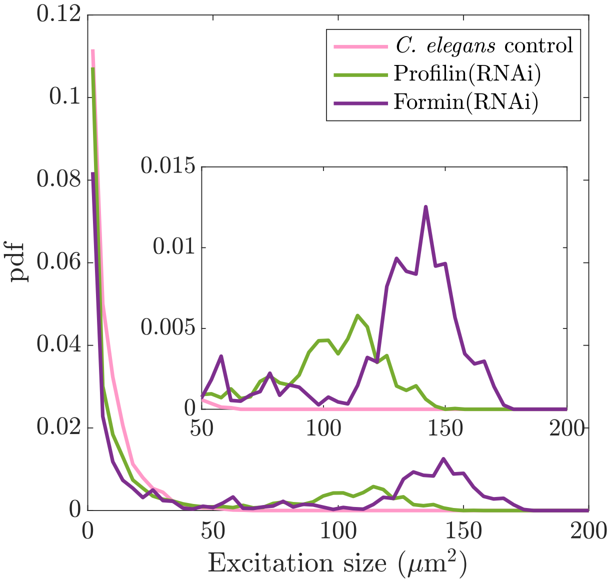

**Figure S3:** PDF of excitation sizes in filtered data across all three experimental conditions. The inset magnifies $x \in [50, 200]$.

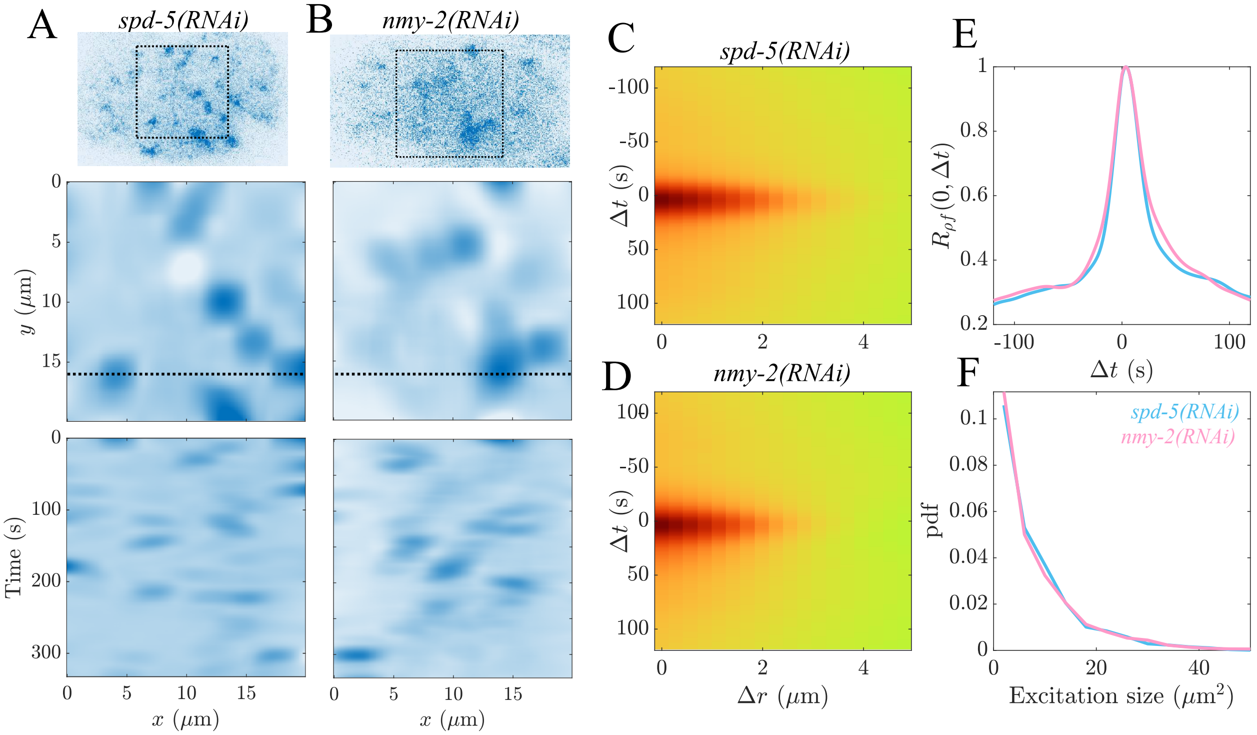

**Figure S4:** Embryos depleted of myosin II manifest the essential features of control embryos. (A,B) Snapshots and kymographs (analogous to those shown in Fig. 1) for *spd-5(RNAi)* embryos (lacking a functional sperm cue for polarization) and *nmy-2(RNAi)* embryos (depleted of myosin II). (C,D) Cross correlation functions $R_{\rho f}$, using colorscale in Fig. 1. (E) Comparing one-dimensional cross correlation along $\Delta r = 0$. (F) Comparing excitation size distributions.

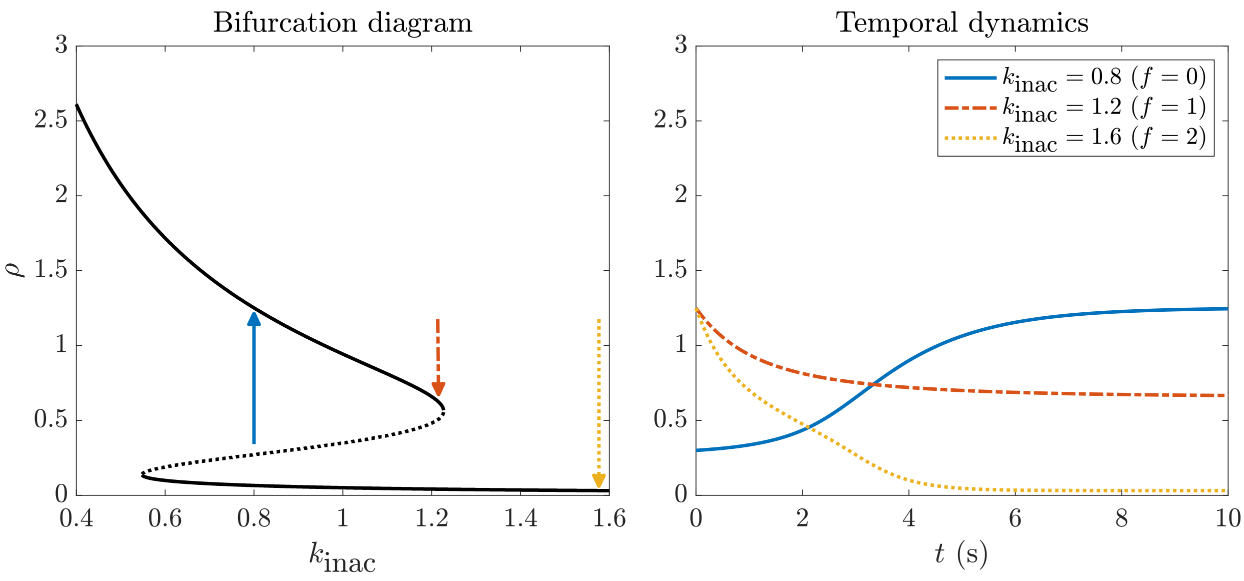

**Figure S5:** Parameterizing the continuum model of Rho. Left: Fixing $k_{\text{b}} = 0.05$, $k_{\text{ac}} = 1$, $K_{\text{sat}} = 0.1$, the steady states of the Rho equation (2a) (with $D_\rho = 0$) produce a bifurcation as a function of $k_{\text{inac}}$. Right: With these parameters, the dynamics of Rho have an approximate intrinsic timescale of less than 5 s. The three scenarios shown correspond to pulse initiation (blue), one actin filament from zero (red), and two actin filaments from zero (yellow).

reduces the steady state equation to a single unknown parameter, the inactivation rate $k_{\text{inac}}$. The bifurcation diagram in Fig. S5 (left panel) shows the steady states of (2a) as a function of $k_{\text{inac}}$. Recalling that $k_{\text{inac}} = k_{\text{inac}}^{(0)} + k_{\text{inh}} f$ in (2c), only $k_{\text{inh}} f$ appears in the inactivation rate, so can fix the effect of a single filament by setting $k_{\text{inh}} = 0.4$. This makes simulations more efficient as only two filaments are sufficient to locally extinguish Rho excitations. Preliminary simulations that vary the basal Rho inhibition parameter $k_{\text{inac}}^{(0)}$ reveal that cross correlations consistent with experimental data can only be generated when the dynamics of Rho are bistable, $0.55 < k_{\text{inac}}^{(0)} < 1.22$ (Section S2.4). The exact value is arbitrary, and we use $k_{\text{inac}}^{(0)} = 0.8$ for the results presented.

The parameters $K_{\text{sat}}$, $k_{\text{b}}$, and $k_{\text{ac}}$ also control the temporal response to step changes in $k_{\text{inac}}$. In Fig. S5, we simulate the response to a decrease/increase in $k_{\text{inac}}$ (which happens when actin filaments are removed/added). We simulate the dynamics of a pulse with no actin filaments ($k_{\text{inac}} = k_{\text{inac}}^{(0)} = 0.8$), and then the dynamics of inhibition with a step change in actin filaments ($k_{\text{inac}} = 1.2$ for one actin filament and $k_{\text{inac}} = 1.6$ for two actin filaments). With the fixed Rho parameters, the Rho dynamics transition to a new steady state over a timescale of about 2–4 s. In practice, since actin subunit lifetimes are about 4 s or more (Fig. 3), this makes the local dynamics of Rho quasi-steady relative to actin.

#### S2.2 Numerical parameters

For the numerical parameters (second section of Table S1), we fix the actin thickness at $g_w = 0.1$ $\mu$m. This is much larger than the thickness of a single actin filament, but when simulating large areas (20 $\mu$m $\times$ 20 $\mu$m), resolving the thickness of a single filament would require $20/0.004 = 5000$ grid cells in each dimension, which would make scanning over parameters impossible. While the thicker radius is primarily set for numerical reasons, it can be interpreted physically as either assuming that the RhoGAP profile is more diffuse than the actin filaments with which it colocalizes, or that

| Parameter | Unit | Definition | Value/range |
|---|---|---|---|
| $k_\mathrm{b}$ | $1/(\mu\mathrm{m}^2{\times}\mathrm{s})$ | Basal Rho activation | 0.05 |
| $k_\mathrm{ac}$ | $1/(\mu\mathrm{m}^2{\times}\mathrm{s})$ | Autocatalysis | 1 |
| $K_\mathrm{sat}$ | $\mu\mathrm{m}^{-6}$ | Saturation | 0.1 |
| $k^{(0)}_\mathrm{inac}$ | 1/s | Basal Rho inactivation | 0.8 |
| $k_\mathrm{inh}$ | $\mu\mathrm{m}^2/\mathrm{s}$ | Actin inhibiting Rho | 0.4 |
| $D$ | $\mu\mathrm{m}^2/\mathrm{s}$ | Rho diffusivity | 0.1 |
| $g_w$ | $\mu$m | Actin thickness | 0.1 |
| $L$ | $\mu$m | Simulated domain size | 20 |
| $\Delta t$ | s | Time step size | 0.25 |
| $\Delta x$ | $\mu$m | Grid spacing | 0.2 |
| $T_\mathrm{fil}$ | s | Actin lifetime | $[0, 40]$ |
| $\nu_p$ | $\mu$m/s | Actin growth/shrinkage rate | $[0, 5]$ |
| $\ell_\mathrm{max}$ | $\mu$m | Max actin length | $[0, 15]$ |
| $q_b$ | $1/(\mu\mathrm{m}^2\times$ s) | Basal nucleation rate | [0,0.3] |
| $\rho^2_\mathrm{max} q_\rho$ | $1/(\mu\mathrm{m}^2\times$ s) | Actin-enhanced nucleation rate | [0,3] |

**Table S1:** Parameters for the hybrid model. The top set of parameters are fixed physical parameters, while the middle set are fixed numerical parameters. The bottom set are varied over the indicated ranges. The maximum steady state of Rho without F-actin ($f = 0$) is denoted by $\rho_\mathrm{max}$.

each "filament" actually represents a bundle of actin filaments (this is our preferred interpretation).

#### S2.3 Estimates from simulated data

Because actin filament assembly dynamics are stochastic, the cross-correlation and excitation statistics vary randomly across individual simulations. Noisy assembly dynamics are amplified by the bistable dynamics of Rho; because excitations cannot form *de novo*, filament levels in some simulations are high enough to quench Rho activity everywhere, resulting in "dead" simulations that we abort before completion. To ensure that we adequately sample each parameter set, we consider a fixed number (ten) of simulations, keeping the first two that last until $t = 240$ s. Parameter sets that cannot sustain at least one simulation out of ten are recorded as being unable to sustain excitations and assigned a low likelihood. Our tests showed that this method estimates the likelihood and prior to within 15%.

#### S2.4 Varying basal inactivation rate

The simulations in the main text all use $k^{(0)}_\mathrm{inac} = 0.8$, so that the excitable dynamics of Rho are stimulated by diffusion, rather than by local depletion of actin (i.e., in the main text, a void in the actin network does not automatically produce an excitation in Rho). In this section, we relax this constraint, allowing $k^{(0)}_\mathrm{inac}$ to vary (with $k_\mathrm{inh} = 0.4$ still fixed) and repeating the parameter estimation in the main text (with a uniform prior).

As in the main text, we generate $5 \times 10^5$ parameter sets, compute the posterior probability of each for the four different data sets, and plot the fifty most likely parameter sets in Fig. S6. The middle and right plots show the same quantities as the main text (Fig. 3). Generally, the trends in these quantities are the same, primarily because the best-fit values for ($k^{(0)}_\mathrm{inac}$) fall into the (bistable) regime used in the main text (right of the dotted line in the left panel). As discussed in Section

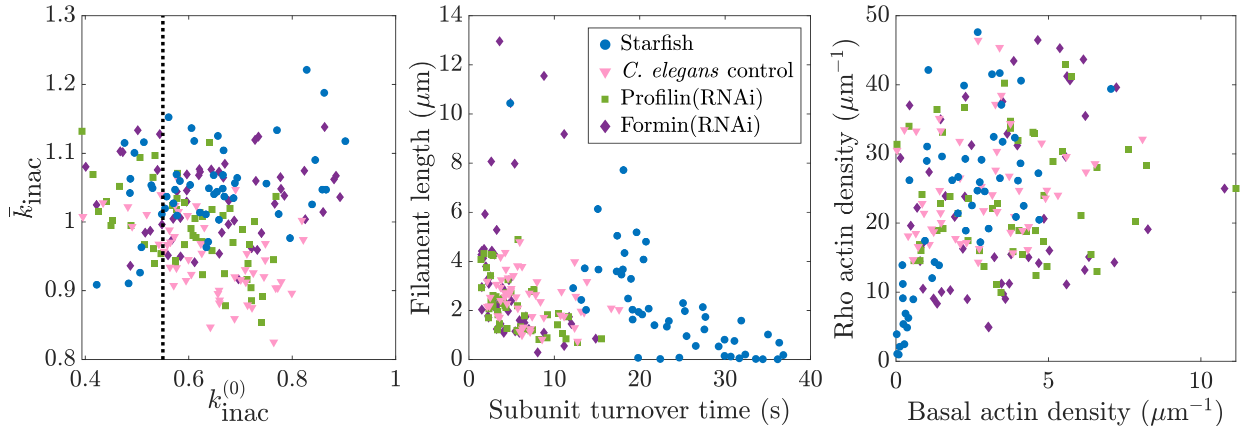

**Figure S6:** Best-fit parameter sets for the case of varied $k_{\text{inac}}^{(0)}$. Similar to Fig. 3 in the main text, we plot the 50 best parameter sets for each data set, this time allowing $k_{\text{inac}}^{(0)}$ to vary. The dotted line shows the boundary of the bistable regime (from the bifurcation plot in Fig. S5). Left: $k_{\text{inac}}^{(0)}$ vs. average inhibition rate $\bar{k}_{\text{inac}} = k_{\text{inac}}^{(0)} + k_{\text{inh}}\bar{f}$. Middle and right: same as main text (Fig. 3)

2.2, this result implies that the the data are best explained by a model in which Rho excitation spreads by diffusion, rather than one in which local depletion of F-actin can induce excitation.

To further understand why the regime of depletion-induced excitation ($k_{\text{inac}}^{(0)} < 0.55$) does not fit the data, we extract the best-fit parameters for the four different data sets with $k_{\text{inac}}^{(0)} < 0.55$, plotting the dynamics in Fig. S7. Aside from significantly more actin in all cases (which is necessary since the basal inactivation rate is lower and more inhibition must come from the filaments), the Rho activity patterns resemble those shown in the main text (cf. Fig. 2), with two important exceptions. First, the best fit to starfish data fails to reproduce the spatial spread of excitations; instead, the dynamics are dominated by local oscillations. This is apparent in the kymographs and in the cross correlation, which lacks the spatial structures of the experimental data and best-fit simulation in Fig. 2. Second, the best fit for *C. elegans* control embryos has a cross correlation which is negative at $(\Delta r, \Delta t) = (0, 0)$, reflecting the tendency for Rho to autoexcite in regions depleted of F-actin. This negative cross-correlation is reduced for best fits to data from profilin- and formin-depleted embryos, in which excitation regions are larger and thus locally positive correlations between Rho and F-actin are more dominant. The failure to match key features of the cross correlations observed for starfish and *C. elegans* controls further supports the conclusion that Rho excitations spread by diffusion, rather than initiating *de novo* in regions of actin voids.

#### S2.5 Unequal growth/shrinkage rates

To examine the assumption of equal growth and shrinkage rates $\nu_p = \nu_d$, we consider the best fit simulations from Fig. 2, for the starfish and *C. elegans* data, with $\nu_p \neq \nu_d$. We set one of $\nu_p$ and $\nu_d$ to the maximum value of 5 $\mu$m/s, then tune the other polymerization speed so that the total time polymerizing and depolymerizing, $T_p = \ell_{\max}/\nu_p + \ell_{\max}/\nu_d$, remains constant. While varying the rates can change the behavior of individual simulations (Fig. S8), the summary statistic of cross correlation is not strongly affected. Notably, there is more qualitative and quantitative change when $\nu_p = 5$ than when $\nu_d = 5$, suggesting that the polymerization rate controls the inferred parameter value when $\nu_p = \nu_d$, while $\nu_d$ is not strongly constrained on its own.

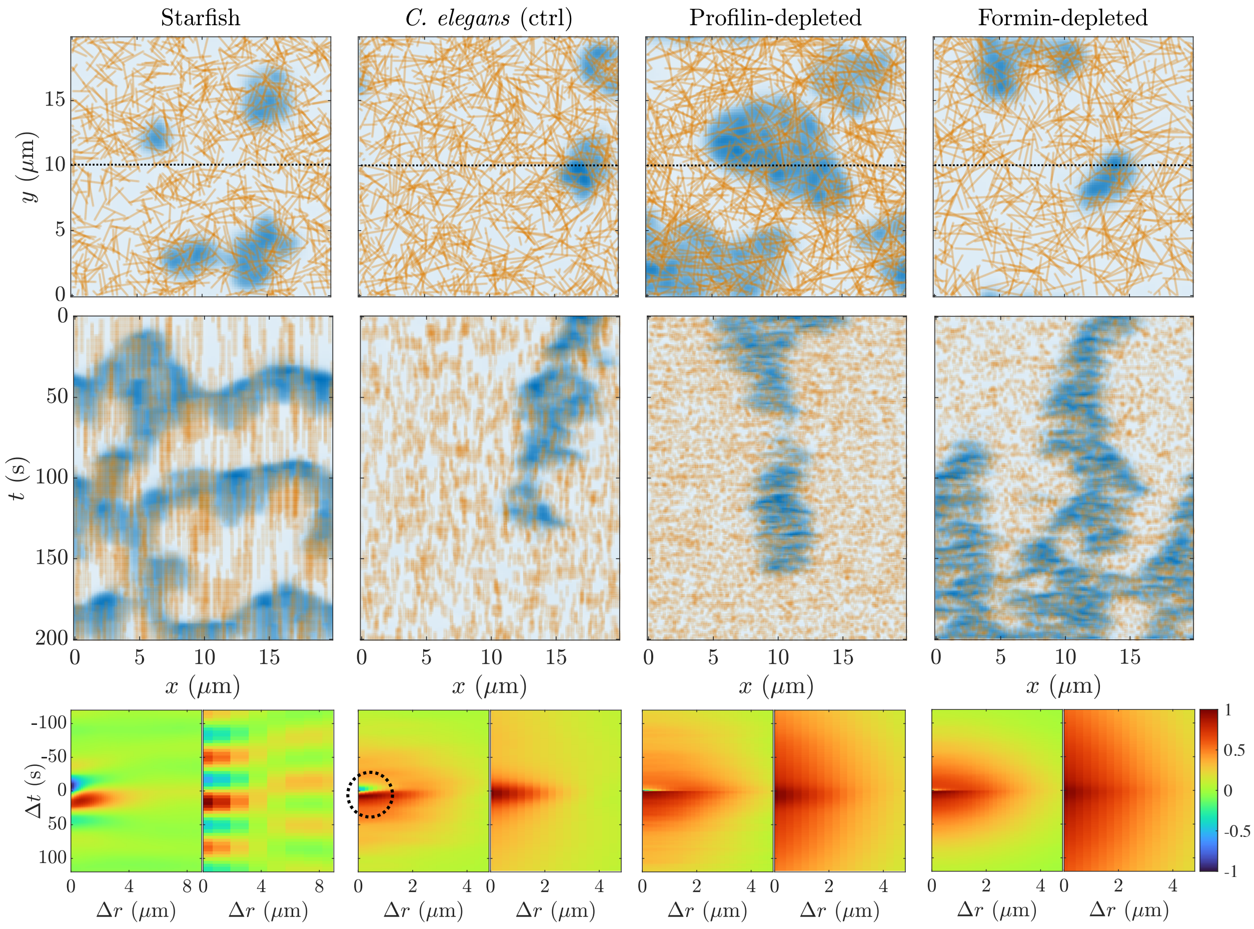

**Figure S7:** Best-fit simulations for each data set assuming $k_{\text{inac}}^{(0)} < 0.55$, i.e., when the dynamics of Rho are automatically excitable without actin. Top: snapshots (at $t = 0$) and kymographs (over $y = 10$). Bottom: cross correlations (left) compared to experimental data (right). Note the (circled) negative cross correlation in the *C. elegans* control parameter set.

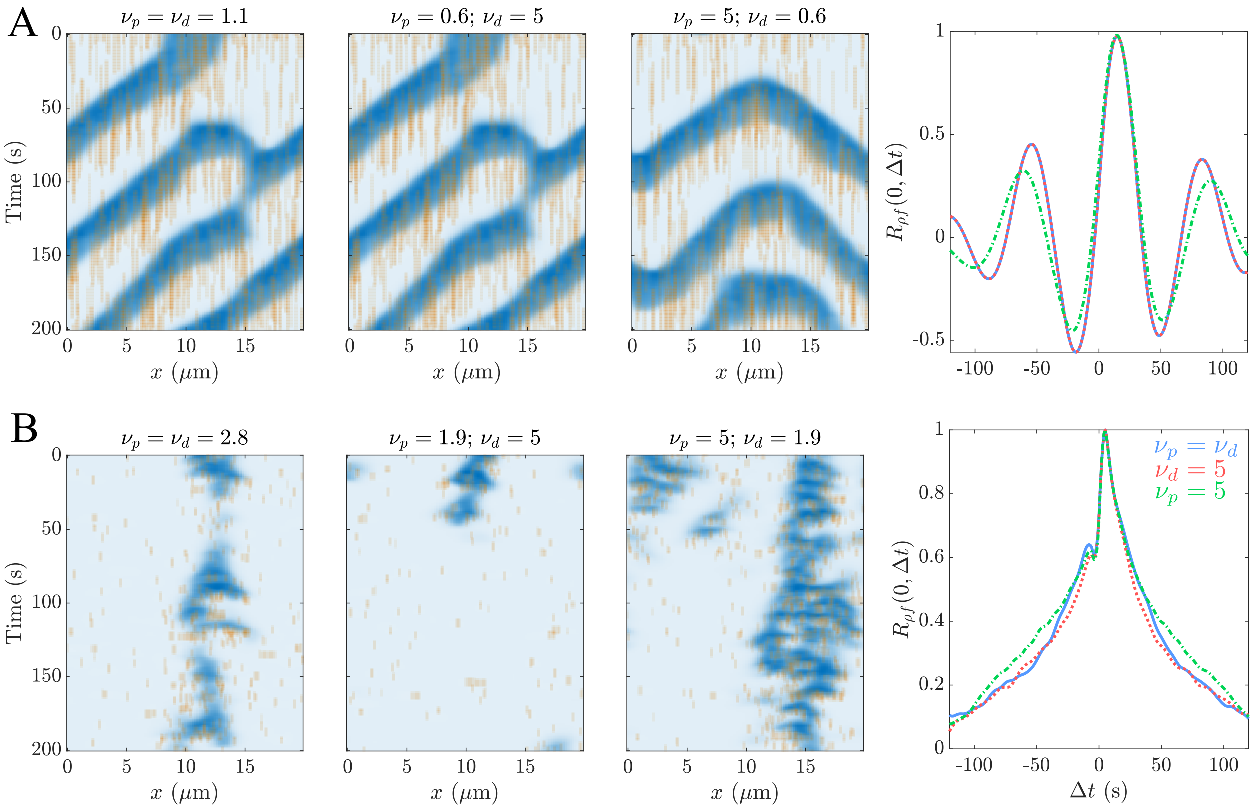

**Figure S8:** Simulations with unequal growth/shrinkage rates. Shown are kymographs of simulations with parameters from Fig. 2 for (A) starfish data and (B) control *C. elegans* data. Kymographs at left show equal growth ($\nu_p$) and shrinkage ($\nu_d$) rates (and match the main text). Middle kymographs show rapid shrinkage rate (5 $\mu$m/s), with growth rates set so that the total filament lifetime is the same as controls. Right kymographs show rapid growth (5 $\mu$m/s), with shrinkage rate set so that the total filament lifetime is the same as controls. Panels at right compare one-dimensional slices of the cross correlations along $\Delta r = 0$.

| Parameter | Starfish | *C. elegans* control | Profilin(RNAi) | Formin(RNAi) |
|---|---|---|---|---|
| $k_{\text{inac}}^{(0)}$ | 0.8,–, 0.538 | 0.8,0.8, 0.479 | 0.8,0.8, 0.428 | 0.8,0.8,0.490 |
| $T_{\text{fil}}$ | 29.4,–, 24.7 | 4.56, 2.30, 5.27 | 4.06, 6.18, 0.432 | 3.61, 5.36, 1.738 |
| $\nu_p$ | 1.13,–, 4.29 | 2.77, 2.15, 1.24 | 3.64, 0.24, 3.51 | 4.44, 2.72, 4.61 |
| $\ell_{\text{max}}$ | 0.0260,–, 1.88 | 2.41, 3.20, 2.66 | 1.18, 1.01, 4.49 | 10.8, 1.33, 3.37 |
| $q_b$ | 0.130, –, 0.069 | 0.00946, 0.100, 0.251 | 0.136, 0.0804, 0.818 | 0.0125, 0.0151, 0.517 |
| $\rho_{\text{max}}^2 q_\rho$ | 1.89, –, 0.530 | 1.28, 0.913, 1.11 | 2.32, 0.899, 1.91 | 0.195, 1.97, 3.99 |

**Table S2:** Parameters for the hybrid model simulations in Figs. 2, 4, and S7. For definitions and units of parameters, see Table S1. Two or three values of each parameter are given; the first is used in Fig. 2, the second in Fig. 4 (omitted for starfish), and the third in Fig. S7.

| Parameter | Fig. S9A | Fig. S9B | Fig. 5B, S10A | Fig. 5C–G, S10B–C |
|---|---|---|---|---|
| $D_\rho$ | 0.1 | 0.1 | 0.1 | 0.1 |
| $k_{\text{b}}$ | 0.05 | 0.05 | 0.05 | 0.05 |
| $k_{\text{ac}}$ | 1 | 1 | 1 | 1 |
| $K_{\text{sat}}$ | 0.1 | 0.1 | 0.1 | 0.1 |
| $k_{\text{inac}}^{(0)}$ | 0.55 | 0.55 | 0.55 | 0.55 |
| $k_{\text{inh}}$ | 0.035 | 0.035 | 0.035 | 0.035 |
| $q_b$ | 0.15 | 0.57 | 0.15 | 0.57 |
| $q_\rho$ | 2 | 7.6 | 2 | 7.6 |
| $k_{\text{diss}}$ | 0.033 | 0.125 | 0.033 | 0.125 |
| $D_f$ | 0.01 | 0.5 | – | – |
| $\nu_p$ | – | – | 0.025 | 0.5 |

**Table S3:** Parameters for continuum modeling in Section 3. Units are given in Table S1.

##### S2.6 Parameter sets

The parameter sets that we use in Figs. 2, 4, and S7 are given in Table S2.

#### S3 Continuum modeling

##### S3.1 Parameter sets

The parameter sets that we use to simulate the PDEs in (2), (S2), and (5a) for Figs. 5, S9, and S10 are given in Table S3.

##### S3.2 Isotropic diffusion

Fig. S9 compares the isotropic directional transport in Fig. 5(A–B) to isotropic diffusion, where the F-actin dynamics are described by

$$\partial_t f = D_f \nabla^2 f + q_b + q_\rho \rho^2 - k_{\text{diss}} f. \tag{S2}$$

As for isotropic advection, slow F-actin diffusivities ($D_f = 0.01$ $\mu$m$^2$/s) yield traveling waves, while fast F-actin diffusivities ($D_f = 0.5$ $\mu$m$^2$/s) yield standing patterns.

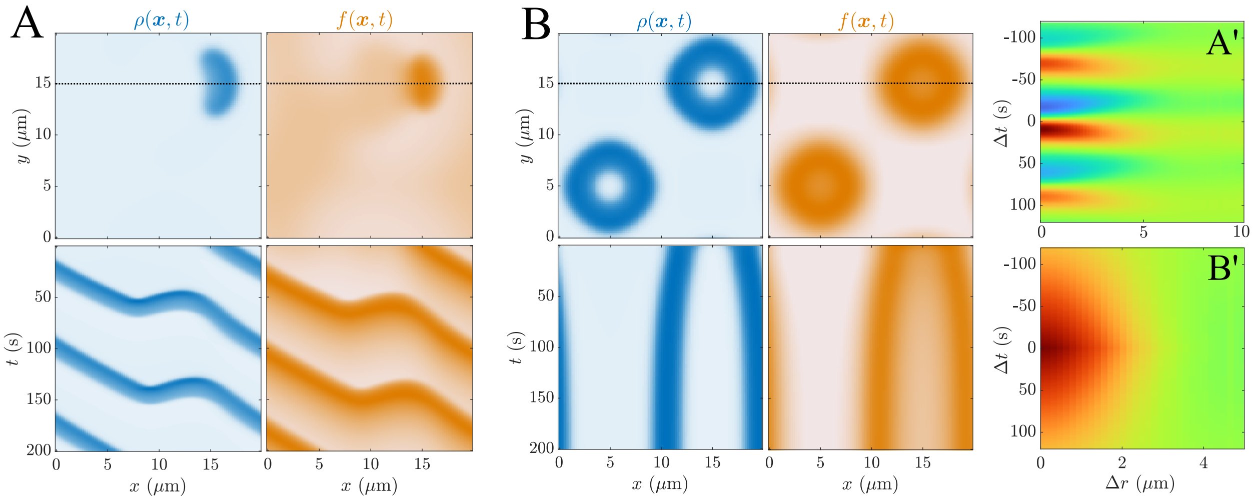

**Figure S9:** Continuum dynamics (2) when F-actin obeys the reaction-diffusion equation (S2) with diffusion rates and turnover times characteristic of (A) starfish and (B) *C. elegans*. Parameters are given in Table S3. The dynamics are similar to those when F-actin evolves by isotropic advection (Fig. 5A–B).

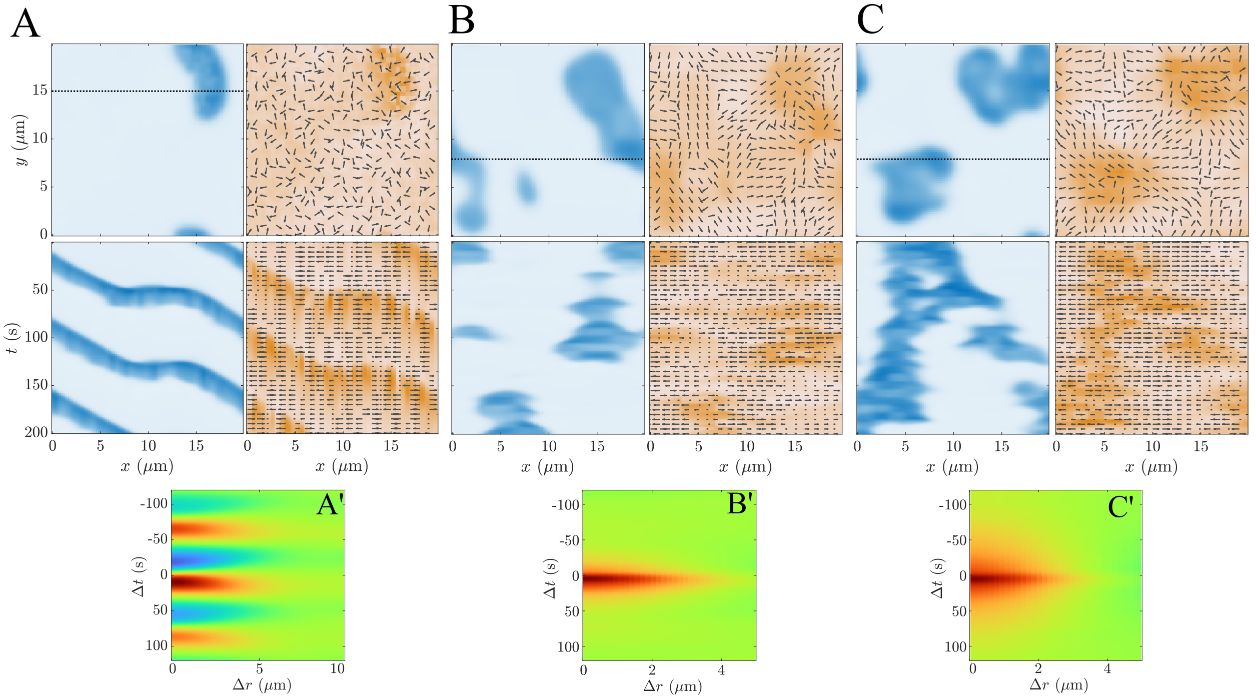

**Figure S10:** Continuum dynamics (2) and (5a) with both spatial and orientational randomness. Using the assembly rates (S3) with eight angles nucleated per region and $c_i \in [0, 2]$, we simulate parameters representing (A) starfish (random region length 0.5 $\mu$m), (B) control *C. elegans* (random region length 3.3 $\mu$m), and (C) profilin/formin depleted *C. elegans* (random region length 1.7 $\mu$m). Other parameters are given in Table S3. (A'-C') Cross correlations.

##### S3.3 Implementing spatial and orientational randomness together

To implement both spatial and orientational randomness, we choose a random sample $\boldsymbol{\Theta}_i$ of $n_\theta$ possible orientations to assemble on each region *and* a region-dependent scaling factor $c_i$, with

$$k_{\mathrm{as}}(\boldsymbol{x} \in \boldsymbol{A}_i, \theta_j) = \begin{cases} c_i 2\pi k_{\mathrm{as}}^{(0)}/n_\theta & \theta_j \in \boldsymbol{\Theta}_i \\ 0 & \theta_j \notin \boldsymbol{\Theta}_i \end{cases}. \quad \text{(S3)}$$

To capture the timescale of filament turnover, we resample $\boldsymbol{\Theta}_i$ and $c_i$ on each region every $1/k_{\mathrm{diss}}$ (values in Table S3). On each region, we nucleate eight out of sixteen angles, and take $c_i \in [0, 2]$.

Fig. S10 shows results of simulations when the region sizes and turnover rates are taken from the hybrid model. To mimic starfish, we set the random region lengthscale to 0.5 $\mu$m and turnover time to 30 s. The small lengthscale makes the F-actin essentially homogeneous, and deterministic dynamics (Fig. 5A) are unaffected. To mimic control *C. elegans* embyos, we set the lengthscale to 3.3 $\mu$m and turnover time to 8 s, similar to the main text. Using both forms of randomness gives a nice mix of transient and mobile excitations, in qualitative agreement with experimental data.

To mimic profilin- and formin-depletion, we simply reduce the region size to 1.7 $\mu$m. This repeats the example in the main text for spatial heterogeneity (Fig. 5E). Using smaller regions with different $\boldsymbol{\Theta}_i$ also gives a more homogeneous distribution of assembly angles on larger scales, similar to the orientational randomness exploration in the main text (Fig. 5G). The increased homogeneity results in larger, more stable excitations, with broader cross correlations.

#### S4 Supplemental movie legends

Video S1 – animated version of unconstrained best-fit simulations in Fig. 2B

- Video S1A – best-fit simulation for starfish data
- Video S1B – best-fit simulation for control *C. elegans* data
- Video S1C – best-fit simulation for profilin-depleted *C. elegans* data
- Video S1D – best-fit simulation for formin-depleted *C. elegans* data

Video S2 – animated version of constrained best-fit simulations in Fig. 4B

- Video S2A – constrained best-fit simulation for control *C. elegans* data
- Video S2B – constrained best-fit simulation for profilin-depleted *C. elegans* data
- Video S2C – constrained best-fit simulation for formin-depleted *C. elegans* data

Video S3 – animated version of continuum simulations in Fig. 5

- Video S3A – isotropic advection with speed corresponding to starfish parameter sets (Fig. 5B)
- Video S3B – isotropic advection with speed corresponding to control *C. elegans* (Fig. 5C)
- Video S3C – random spatial actin nucleation with lengthscale 3.3 $\mu$m (Fig. 5D)
- Video S3D – random spatial actin nucleation with lengthscale 1.7 $\mu$m (Fig. 5E)
- Video S3E – random orientational actin nucleation with one nucleated angle (Fig. 5F)
- Video S3F – random orientational actin nucleation with six nucleated angles (Fig. 5G)

Video S4 – animated version of best-fit simulations in Fig. S7 with $k_{\mathrm{inac}}^{(0)} < 0.55$ (the depletion-induced excitability regime)

- Video S4A – best-fit simulation for starfish data when $k_{\mathrm{inac}}^{(0)} < 0.55$

- Video S4B – best-fit simulation to control *C. elegans* data when $k_{\text{inac}}^{(0)} < 0.55$
- Video S4C – best-fit simulation for profilin-depleted *C. elegans* data when $k_{\text{inac}}^{(0)} < 0.55$
- Video S4D – best-fit simulation for formin-depleted *C. elegans* data when $k_{\text{inac}}^{(0)} < 0.55$

Video S5 – animated version of simulations with both positional and orientational and randomness (Fig. S10)

- Video S5A – simulation mimicking starfish data (Fig. S10A)
- Video S5B – simulation mimicking control *C. elegans* data (Fig. S10B)
- Video S5C – simulation mimicking profilin- and formin- depleted *C. elegans* (Fig. S10C)
